# Explainable AI and Multiclassifiers for Staging Biomarker Discovery in Lung Squamous Cell Carcinoma

**DOI:** 10.1101/2025.07.10.663226

**Authors:** Débora V. C. Lima, Patrick Terrematte, Beatriz Stransky, Adrião D. D. Neto

## Abstract

Lung cancer is one of the most common and lethal types of cancer worldwide. Among its subtypes, lung squamous cell carcinoma (LUSC) is one of the most frequent. Identifying biomarkers for LUSC represents a significant challenge due to its high molecular heterogeneity. However, this promising search may elucidate biological mechanisms and reveal potential therapeutic targets. In this context, the present study used gene expression data from TCGA-LUSC, combined with feature selection, data balancing, machine learning, and explainable artificial intelligence (XAI) to identify possible biomarkers related to staging. The employed methods demonstrated robust classification metrics and highlighting random forest, which achieved an accuracy of 0.91. The use of data balancing and feature selection techniques proved to be crucial in the classification process. In addition, it was possible to identify the 16 most relevant genes selected by random forest using the SHapley Additive Explanations (SHAP) method. Among them, three genes (MYOSLID, IMPDH1P8, and COL9A3) were chosen by all successful classifiers, positioning themselves as potential staging biomarkers and possible molecular therapeutic targets for LUSC.

## Introduction

Lung cancer is one of the most commonly diagnosed types of cancer around the world in men and women. Its incidence levels vary according to the population profile, such as geographic location, cigarette smoking, and the patient’s gender, generally being more common in men [1]. Lung cancer has high incidence and mortality rates; in addition, there were delays in diagnosis, treatments, and recovery of patients during the COVID-19 pandemic, leading to an increase in the disease in advanced stages and mortality [2].

The most common lung cancer is lung adenocarcinoma, followed by lung squamous cell carcinoma (LUSC). While lung adenocarcinoma has already benefited from molecular treatments, LUSC still lacks effective molecular targets [3]. This occurs because the molecular changes which define lung adenocarcinomas and LUSC are distinct [4].

Developing molecularly targeted therapies for LUSCs presents significant challenges, leaving patients in advanced stages with few treatment options. This is problematic since the majority of patients are diagnosed in advanced stages and have high mortality associated with the disease [5]. However, studies of large projects such as TCGA-LUSC are a promising strategy for discovering possible druggable molecular biomarkers [6]. According to [7], the most informative features for the prognosis of LUSCs are methylation, copy number alterations, and particularly gene expression.

Considering the problem of searching for biomarkers in LUSCs and the availability of a large volume of molecular data, it is necessary to use techniques capable of dealing with complex data [8]. In this context, machine learning techniques emerge as alternatives for discovering relevant information from molecular data [9].

The use of machine learning and artificial intelligence (AI) techniques is promising. Machine learning and bioinformatics associated with cancer data can be used in several applications, such as drug discovery, survival prediction, prognosis prediction, diagnosis, image analysis, and the search for biomarkers [10], [11], [12], [13], [14], [15], which has the potential to make medicine increasingly personalized. However, as the complexity of the data and the problem increases, the less interpretable the models become. In turn, the field of explainable artificial intelligence (XAI) is advancing to make AI models more transparent and interpretable [16].

Shapley Additive Explanations (SHAP) is a model-agnostic XAI approach which aims to provide insights into the inner workings of complex models and quantitative visualizations of model predictions, which is essential in AI models for biological problems [17]. The use of XAI applied to lung cancer has already been used in different applications, such as for predicting EGFR mutation [18], predicting length of hospital stay [19], diagnosis [20], discovery of biomarkers [21], toxicity prediction [22], and survival prediction [23].

Therefore, the present work aims to use machine learning and XAI techniques associated with gene expression data to obtain a staging-associated signature for LUSC. This signature can be used as a staging biomarker and studied as a possible therapeutic target, enabling discovery of new information about LUSC and increasingly personalized medicine.

## Material and Methods

Fig. 1 summarizes the methodology from data recovery to obtaining the gene signature.

**Figure 1.**
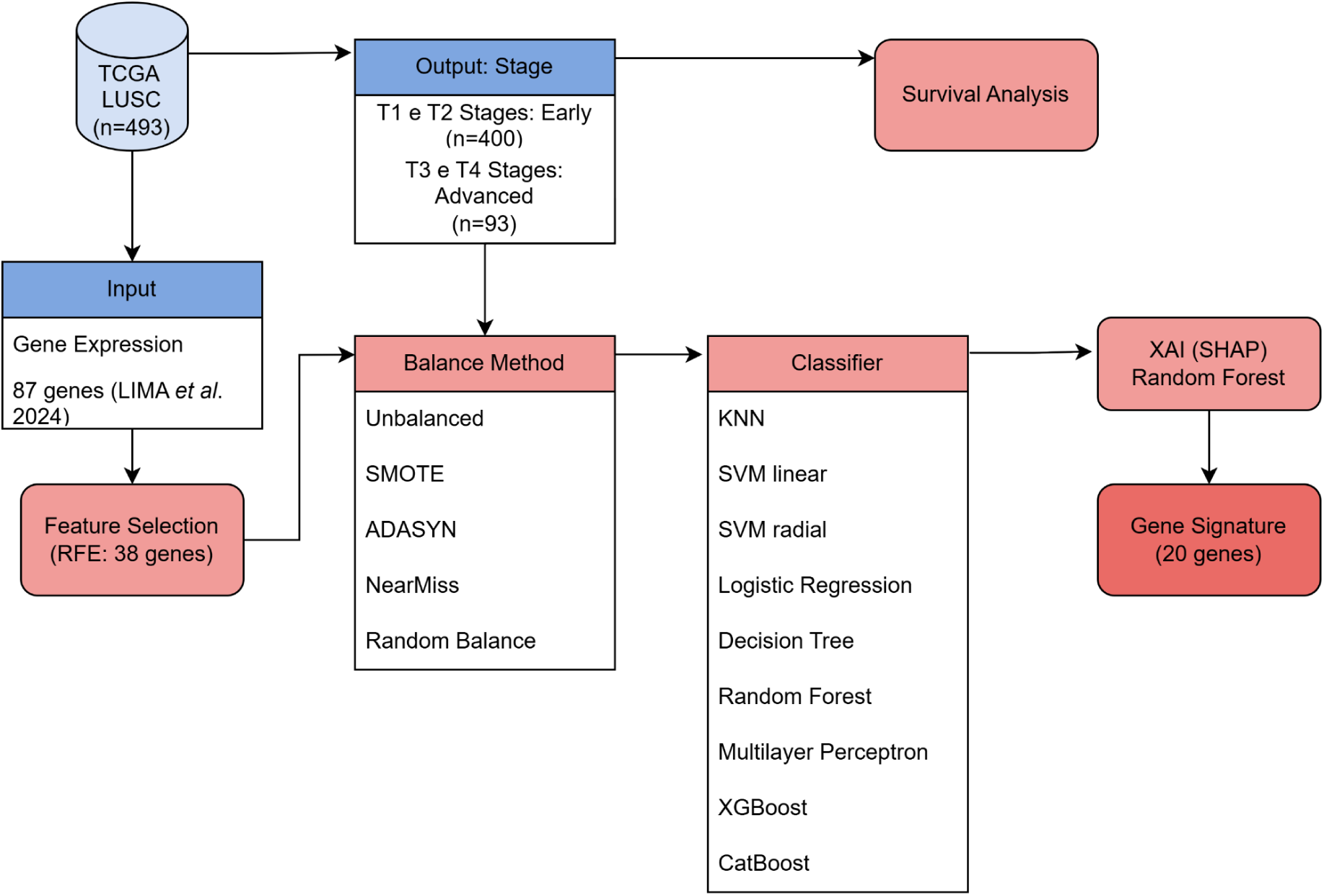
Workflow including data retrieval and processing, feature selection with recursive feature elimination, balance methods, machine learning methods for classification, explainable artificial intelligence with Shapley Additive Explanations, and the resulting gene signature.

### Data Exploration and Pre-Processing

The clinical and gene expression data (count data and log+1 normalization were used) of 493 patients (without normal samples and patients with missing clinical data) were obtained from the LUSC TCGA project in the GDC Data Portal [24]. Their data were retrieved using the TCGABiolinks package (2.24.3) [25] and Xenabrowser [26].

Clinical data were processed with the Pandas library (version 2.2.3) [27], and patient staging was selected as the output variable. The staging was divided into two binary categories for classification: patients T1 and T2 (T1 and T2 = 0) were considered early stage (n = 400), and patients T3 and T4 (T3 and T4 = 1) were regarded as advanced stage (n = 93). A survival analysis was additionally performed to compare early vs. advanced stages using the Lifelines library (version 0.30.0) [28]. We used normalized gene expression data (logarithmic normalization) of 87 genes associated with survival in LUSC for input data which had been selected in a previous study [29].

### Feature Selection

A recursive feature elimination (RFE) was performed as the feature selection method using the Sklearn library (version 1.5.2) [30]. RFE used logistic regression as hyperparameters, cross-validation (stratified kfold = 5) as an estimator, and the area under the curve (ROC_AUC) as scoring to evaluate the best model results.

### Data Balancing

Tests were conducted using unbalanced data and four balancing methods. The unbalanced data referred to the 493 patients recovered from the TCGA-LUSC project. In turn, three methods for the balancing process were used from the imbalanced-learn library (version 0.12.4) [31]. Two of these methods were oversampling (SMOTE and ADASYN), and one method was undersampling (NearMiss). Furthermore, another balancing method was performed with undersampling, randomly keeping only half of the patients in the majority class.

### Machine Learning Methods

We initially tested nine methods to classify the staging of individuals to classify them. The k-nearest neighbors (KNN), radial and linear support vector machine (SVM), logistic regression (LR), decision tree (DT), random forest (RF), and multilayer perceptron (MLP) algorithms were applied using the Sklearn library (version 1.5.2) [30]. The XGBoost library (version 2.1.3) [32] and the CatBoost library (version 1.2.7) were used [32].

Each method was tested with 87 input genes and the genes were selected by RFE as input. In addition, all methods were tested with and without cross-validation. When cross-validation (CV) was applied, 5 folds were used; when cross-validation was not applied, the validation of the dataset was divided into proportions of 70% - 30% for the training and testing groups, respectively.

### Evaluation of Model Performance

Next, we used the following metrics to evaluate all of the models’ performance: area under the curve (AUC), accuracy, F1 score, precision, sensitivity, and specificity. The metrics and ROC curves with model performance were obtained using the Sklearn library (version 1.5.2) [30] and Matplotlib (version 3.9.2) [33]. We conducted a systematic search for literature-based gene signatures related to LUSC to compare our proposed gene signature with existing models. Then, we applied SMOTE for class balancing and used stratified 5-fold cross-validation during model training to ensure sound and unbiased evaluation, consistent with the previous steps of our pipeline. We compared our full 38-gene signature and a 20-gene subset based on feature importance against 11 literature-based gene signatures using a set of six machine learning models, including: CATBoost, XGBoost, random forest, support vector machines (with both radial basis function and linear kernels) and logistic regression. The inclusion criteria required that studies be published from 2022 onward and incorporate RNA-Seq data from the TCGA-LUSC project as one of the primary data sources. Only studies that presented gene signatures or biomarkers specifically developed for LUSC classification, prognosis, or treatment response were considered.

### Explainable Artificial Intelligence

We employed Shapley Additive Explanations (SHAP) (version 0.46.0) [34] for explainable artificial intelligence (XAI) to generate a summary plot with the 20 top genes. This plot allows us to evaluate the impact of the features in the validation set based on the classifications performed.

The summary plots of SHAP values were plotted for all classifiers. When cross-validation was applied, the plot was generated for each fold to verify the consistency in selecting the most impactful genes.

### Diagrams and Upset Plots

Next, we created Venn diagrams and Upset plots using the Venn (version 0.1.3), Matplotlib (version 3.9.2) [33], Matplotlib-venn (version 1.1), and upsetplot (version 0.9.0) [35] libraries to demonstrate the intersections of genes selected by different classifiers. The intersections of the gene signatures obtained with the best performances were verified (the top 20 most impactful genes for classification selected by SHAP). Furthermore, the genes selected by the folds when cross-validation was applied were also compared.

### Pathway Fluxograms

The KEGG database was used to obtain information about the participation of signature genes in pathways associated with lung cancer genes [36], [37], [38]. A summarized version of the pathway was used to generate the flowcharts, with a focus on the processes associated with the genes of interest in PathwayMapper [39].

### Development

We developed all scripts using Python (version 3.10) [40] on Google Colab [42] and using the R programming language (version 4.1) [41] on RStudio [42]. The implementations in R were conducted on the RStudio server workstation of BioMe (Bioinformatics Multidisciplinary Environment) at Metropolis Digital Institute (IMD) from the Federal University of Rio Grande do Norte. The scripts are available on GitHub (https://github.com/Debora96/ArticleXAI).

## Results

### Input and Output Data

We performed recursive feature elimination (RFE) to reduce the number of input features, which enabled reducing the input from 87 genes to 38 genes. We subsequently used these 38 genes as input in the algorithms tested for the staging classification (Table 1).

**TABLE I.**
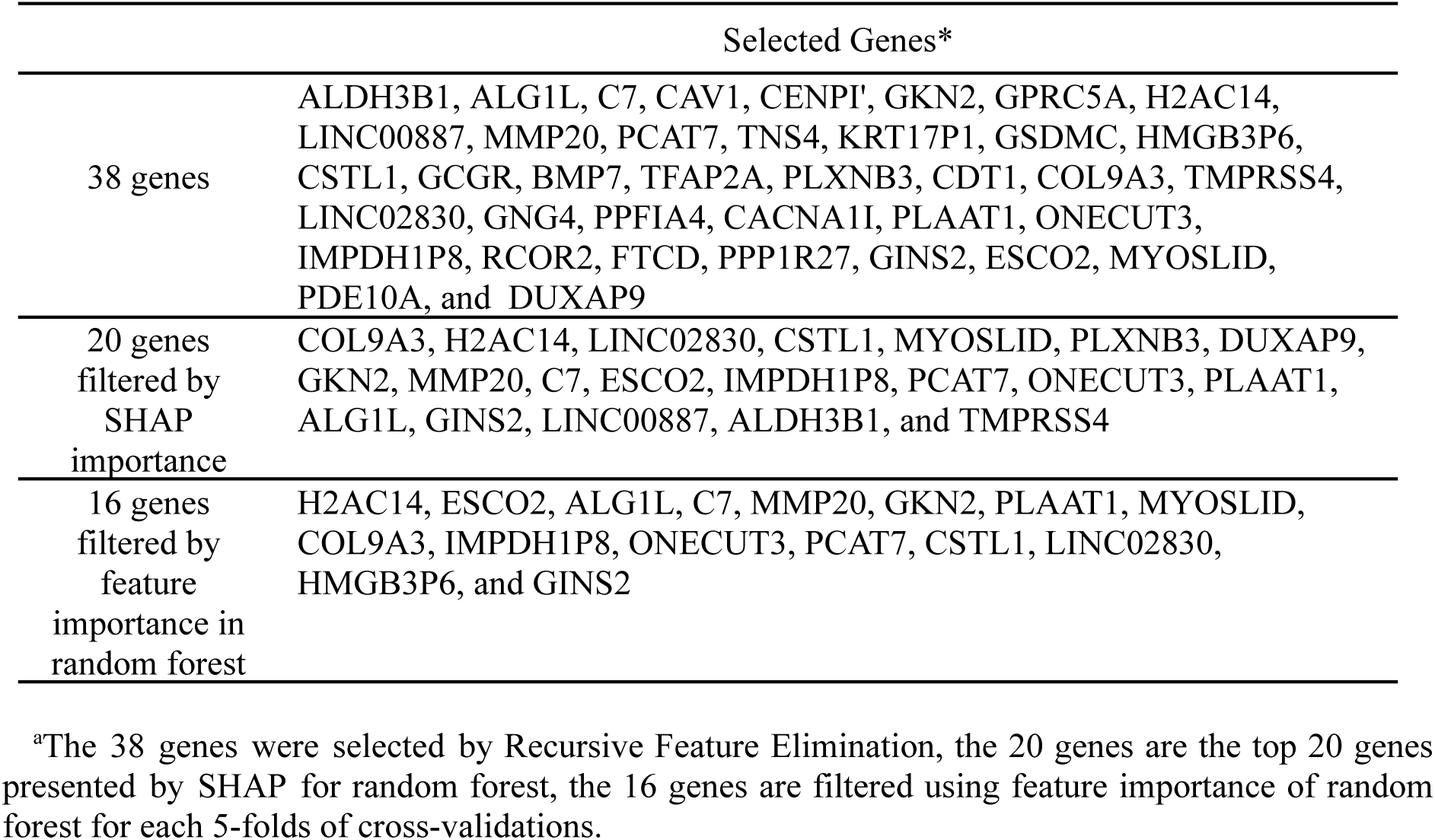
Genes Selected by Explainable AI and Filtered by Multiclassifiers.

Then, we used tumor staging for the output variable to classify patients into early and late stages. We also conducted a survival analysis considering the two output groups, and (as expected) we confirmed that patients in more advanced stages have lower survival rates compared to patients in the early stages (log-rank p-value = 0.00197) (Supplementary Fig. 1).

Furthermore, we had groups with different numbers of individuals due to the use of balancing techniques, as well as different proportions between the early and late-stage classes (Table 2).

**TABLE II.**
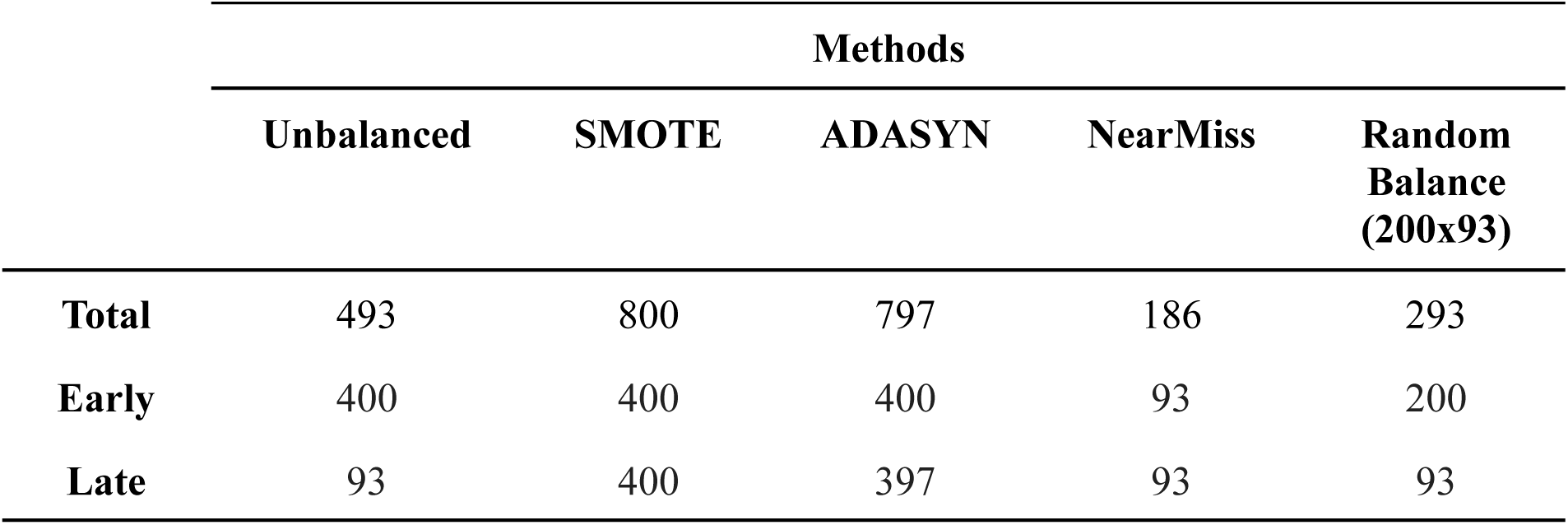
Genes Selected by Explainable AI and Filtered by Multiclassifiers.

### Staging Classification

We performed staging classification with different methods. Tests were performed for all classification techniques, with 87 input genes and also with 38 genes selected by RFE. In addition, cross-validation tests were also conducted in both cases to increase the reliability of the results. As a result, we observed that the best results were using 38 genes from RFE and cross-validation.

Oversampling methods generated data to approximate the number of individuals from the minority class to the majority class. In contrast, undersampling methods selected data to approximate the number of individuals from the majority class to the minority class. As can be seen in Table 3, tests using RFE and cross-validation with unbalanced data and using undersampling techniques showed poor performance. The NearMiss technique was the under-sampling technique with the best results. However, the best average F1 scores for NearMiss were between 0.61 and 0.75.

**TABLE III.** Performance of Classification Techniques Using Recursive Feature Elimination, Cross-Validation and Different Balancing Methods.

| **Mean F1 Score** |  |  |  |  |  |
| --- | --- | --- | --- | --- | --- |
| **Method** | **Over-sampling** |  |  | **Under-Sampling** |  |
|  | **Unbalanced** | **SMOTE** | **ADASYN** | **Near Miss** | **Random Balance\***<br>**(200x93)** |
| **MLP** | 0.0 | 0.36 | 0.67 | 0.64 | 0.0 |
| **KNN** | 0.05 | 0.76 | 0.73 | 0.36 | 0.23 |
| **SVM linear** | 0.30 | 0.79 | 0.76 | 0.53 | 0.43 |
| **SVM radial** | 0.0 | 0.80 | 0.77 | 0.63 | 0.0 |
| **LR** | 0.32 | 0.78 | 0.75 | 0.75 | 0.43 |
| **DT** | 0.20 | 0.76 | 0.71 | 0.55 | 0.36 |
| **XGBoost** | 0.13 | 0.88 | 0.89 | 0.59 | 0.30 |
| **CatBoost** | 0.05 | 0.90 | 0.91 | 0.59 | 0.24 |
| **RF** | 0.04 | 0.91 | 0.90 | 0.63 | 0.12 |

<sup>a</sup>The random balance technique consisted of the random selection of half of the majority class as an undersampling strategy.

Both SMOTE and ADASYN had satisfactory performances for over-sampling techniques (average F1 Scores) for almost all classification techniques. Except for multilayer perceptrons, oversampling techniques had performances between 0.72 and 0.92. We can highlight SMOTE balancing and the XGBOOST, CatBoost, and random forest classification methods as the best-performing methods (Fig. 2).

**Figure 2.**
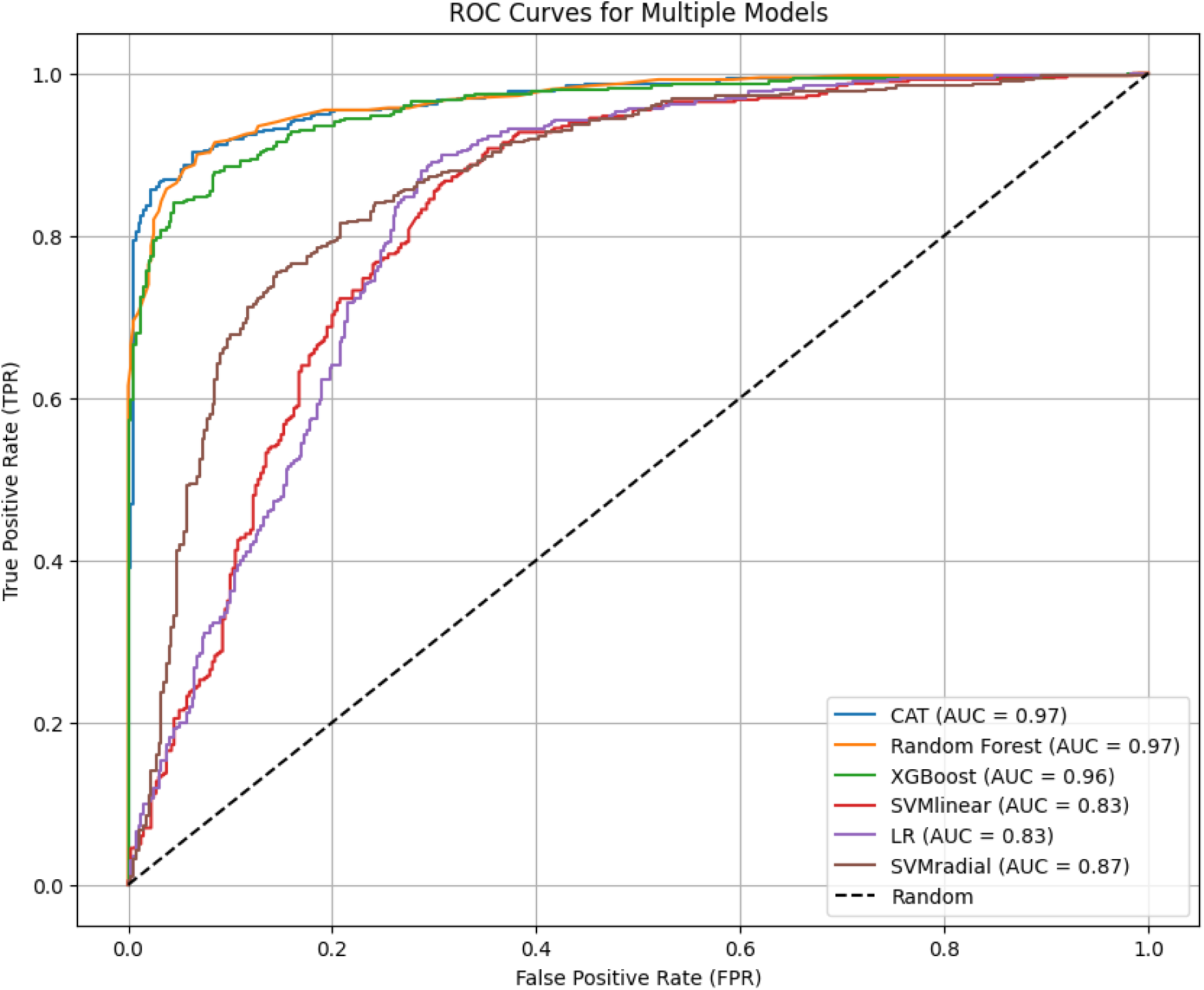
ROC Curves for multiple models using the SMOTE balance method.

Among the best classification methods, CatBoost and Random Forest had all metrics (accuracy, F1 score, precision, sensitivity, and specificity) between 0.89 and 0.92; they also presented very close AUCs (0.973 for Random Forest and 0.975 for CatBoost), presenting practically similar results (Table 4).

**TABLE IV.**
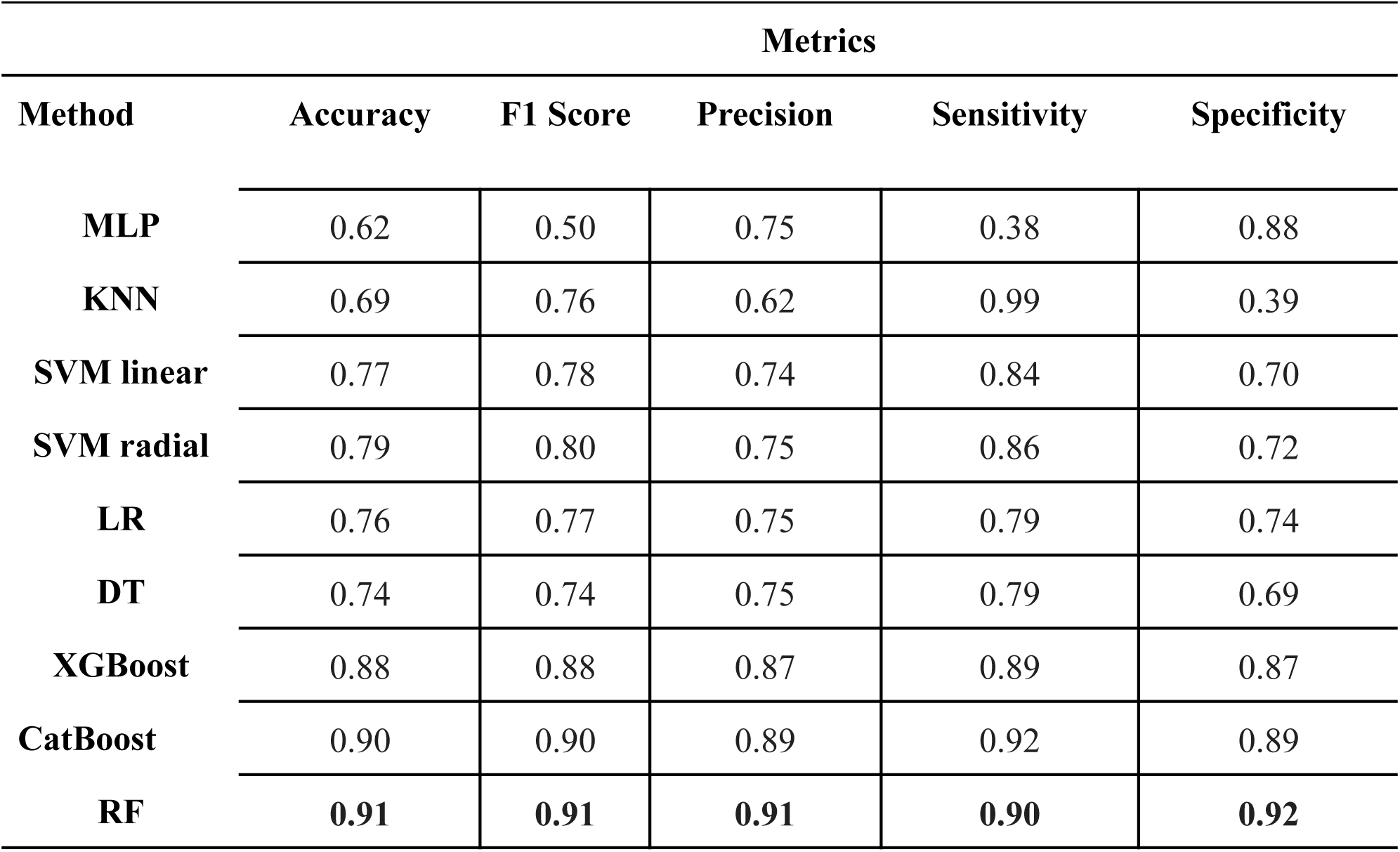
Methics of Classification Techniques Using Recursive Feature Elimination, Cross-Validation and SMOTE.

| Metrics |  |  |  |  |  |
| --- | --- | --- | --- | --- | --- |
| Method | Accuracy | F1 Score | Precision | Sensitivity | Specificity |
| MLP | 0.62 | 0.50 | 0.75 | 0.38 | 0.88 |
| KNN | 0.69 | 0.76 | 0.62 | 0.99 | 0.39 |
| SVM linear | 0.77 | 0.78 | 0.74 | 0.84 | 0.70 |
| SVM radial | 0.79 | 0.80 | 0.75 | 0.86 | 0.72 |
| LR | 0.76 | 0.77 | 0.75 | 0.79 | 0.74 |
| DT | 0.74 | 0.74 | 0.75 | 0.79 | 0.69 |
| XGBoost | 0.88 | 0.88 | 0.87 | 0.89 | 0.87 |
| CatBoost | 0.90 | 0.90 | 0.89 | 0.92 | 0.89 |
| RF | **0.91** | **0.91** | **0.91** | **0.90** | **0.92** |

Considering the interpretability, performance, and wide use of this technique for biomarker selection problems, we chose to continue using Random Forest for explainable artificial intelligence analyses.

### Explainable Artificial Intelligence and the Impact of Features

The SHAP method was used in the explainable AI analysis to verify which of the 38 genes were most important in the classification process performed by random forest.

Fig. 3 shows the top 20 selected genes in each of the five folds of the random forest classification. From this analysis, we observed that the COL9A3 gene was the most important in three of the five folds, and the GKN2 gene was the most important in two folds. Furthermore, among the 20 genes selected by each fold, all five folds shared 16 genes.

**Figure 3.**
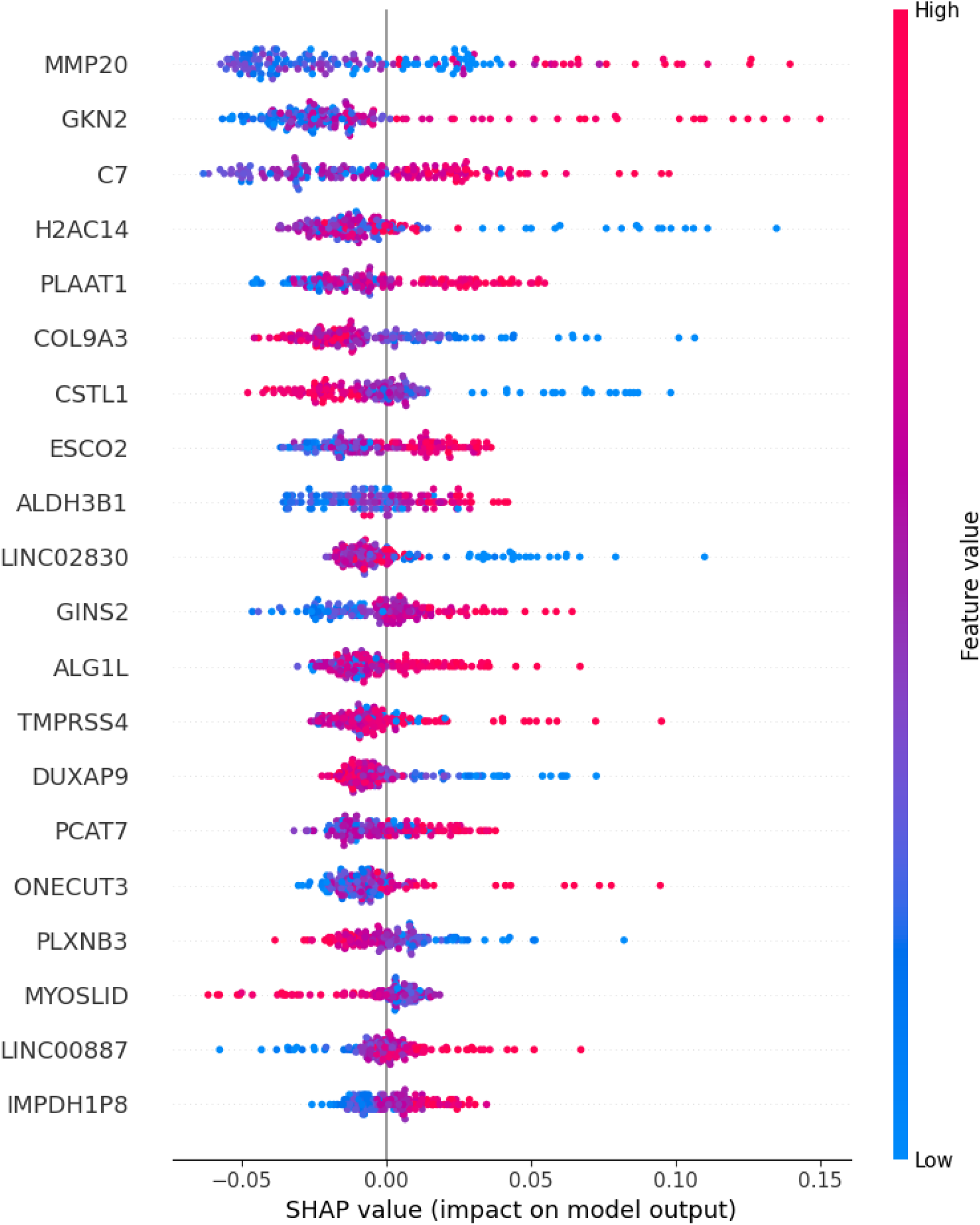
SHAP summary plot with the distribution of the SHAP values for each feature. This summary plot presents a random forest analysis and their top 20 genes. Each dot represents a SHAP value for a feature per patient, and the color represents the feature value from high to low, respectively.

In addition to the good performance of random forest, cross-validation enabled us to observe that the top genes which impacted the classification in the five folds were quite similar. In turn, we evaluated the contributions of variables in the decision-making process. Advanced staging (Fig. 3) was associated with lower expression values in the COL9A3, H2AC14, LINC02830, CSTL1, MYOSLID, PLXNB3, and DUXAP9 genes, and a higher expression of GKN2, MMP20, C7, ESCO2, IMPDH1P8, PCAT7, ONECUT3, PLAAT1, ALG1L, GINS2, LINC00887, ALDH3B1, and TMPRSS4 genes.

### Benchmarking Literature-Based Gene Signatures for Lusc Staging Classification

We initially conducted an analysis with gene signatures previously reported in 11 gene signatures on the literature reported in recent LUSC studies aiming to compare the performance of our proposed 38-gene signature for LUSC staging (Supplementary Table 1). The signatures used for comparison were generated by regression or correlation techniques, in which we evaluated and compared signatures from other studies with the same algorithms tested in our signature to verify how each one performed in classifying patient staging. Fig. 4 compares the literature-based gene signatures to a set of machine learning models used in this study.

**Figure 4.**
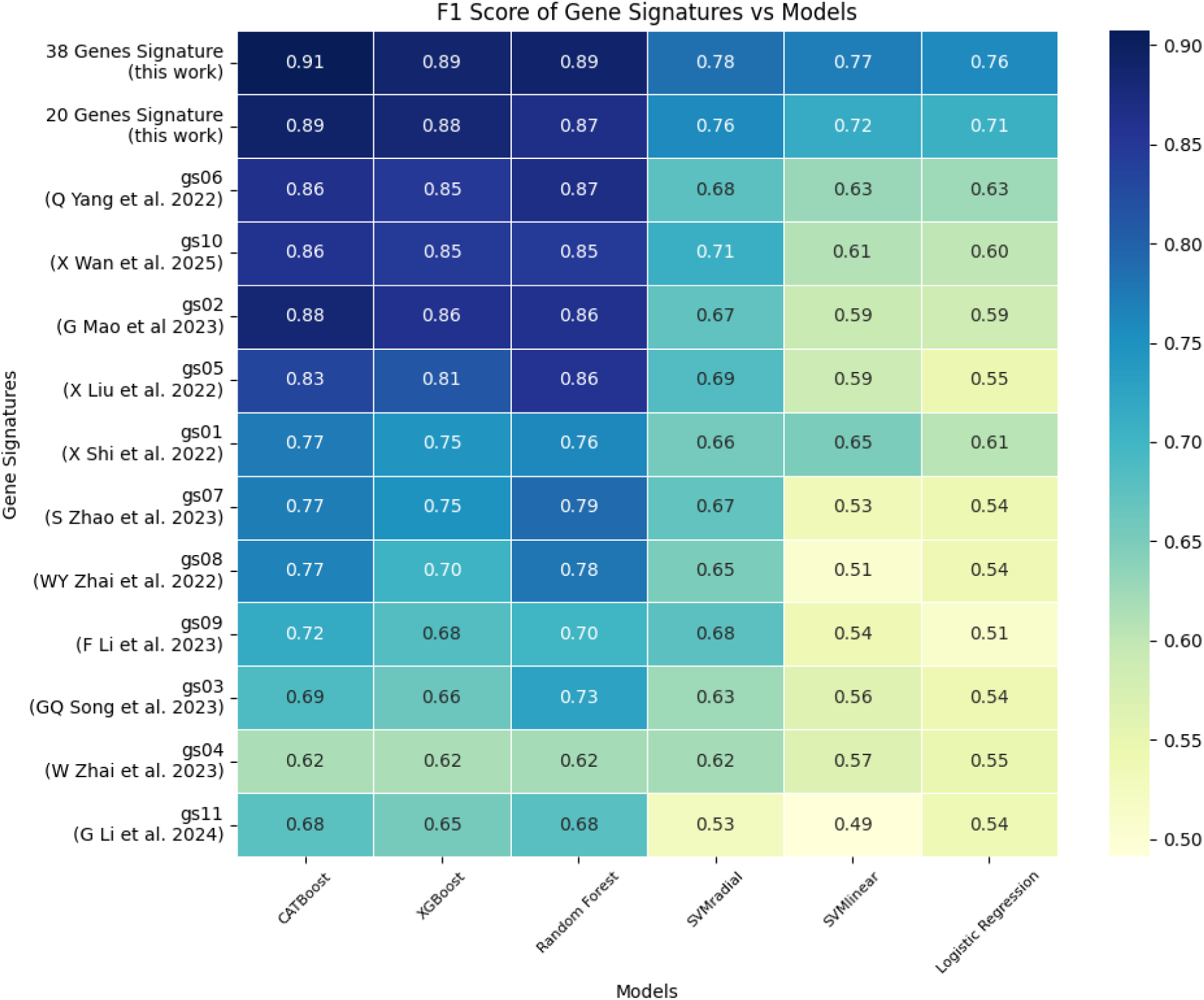
Benchmarking literature-based results with the 38-gene signature evaluated using F1 score for CATBoost, XGBoost, random forest, support vector machine with radial basis function and linear kernels, and logistic regression models.

As a result, the CATBoost model achieved the highest performance with a F1 score of 0.91, indicating discriminative capability in classifying LUSC samples based on the 38-gene signature.

### The Novel 38-Gene Signature

After observing the performance of the random forest with cross-validation in association with SHAP, we obtained the top 20 genes for each of the five folds. As can be seen in Fig. 5, these groups had 16 genes in common (H2AC14, ESCO2, ALG1L, C7, MMP20, GKN2, PLAAT1, MYOSLID, COL9A3, IMPDH1P8, ONECUT3, PCAT7, CSTL1, LINC02830, HMGB3P6, and GINS2).

**Figure 5.**
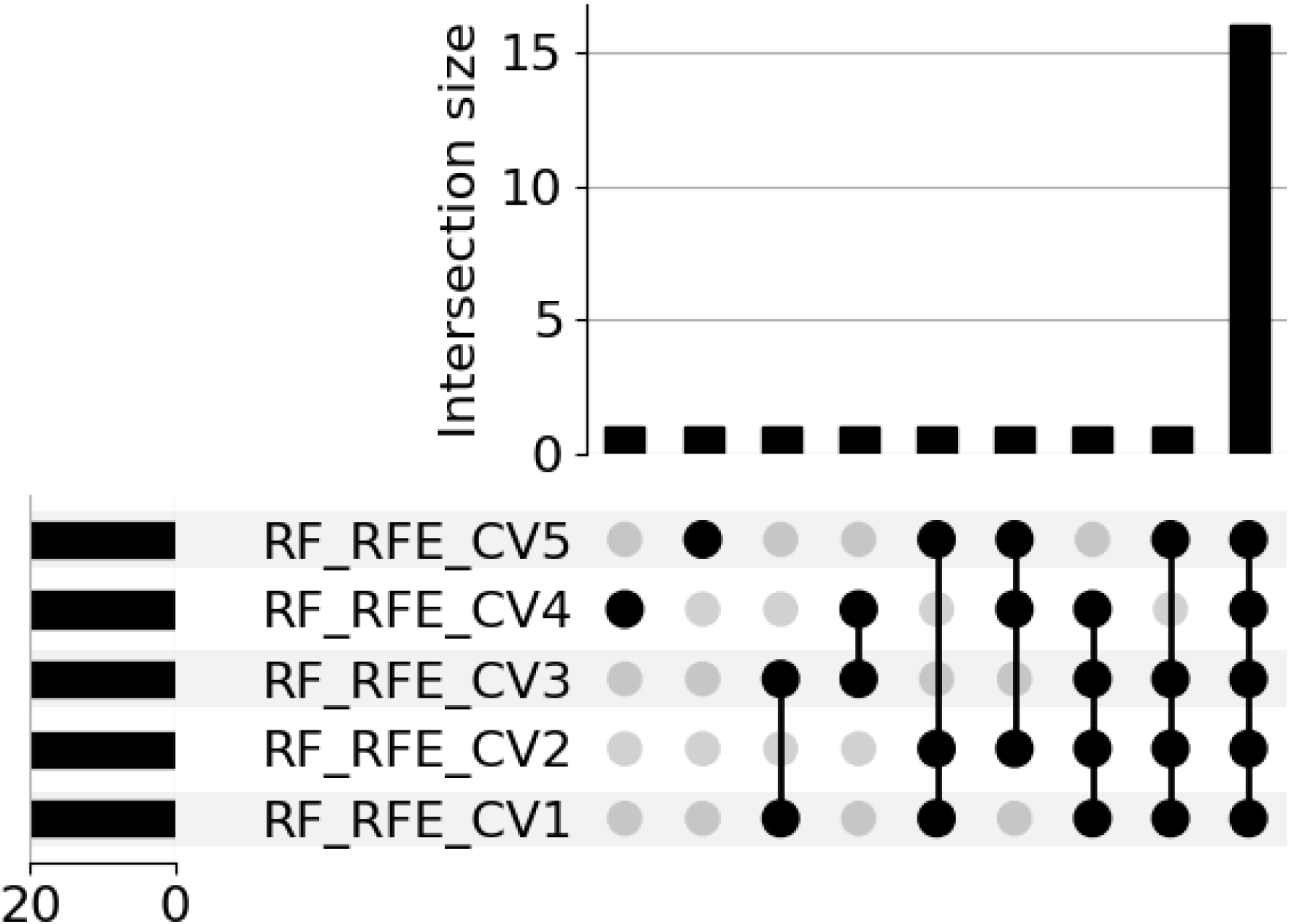
Upset plot with intersections of the fi ve folds from random forest. In addition to the 16 genes selected in all folds, another eight of the genes were selected from the folds. Two genes were unique to just 1 fold, CACNA1I (fold 4) and DUXAP9 (fold 5). The ALDH3B1 gene was selected by folds 1 and 3, and the FTCD gene was selected by folds 3 and 4. The GCGR gene was selected by folds 1, 2, and 5, and the TMPRSS4 gene was selected by folds 2, 4, and 5. The GSDMC gene was selected by folds 1, 2, 3, and 4, and finally, the LINC00887 gene was selected by folds 1, 2, 3, and 5.

Furthermore, we observed 3 genes (MYOSLID, IMPDH1P8, and COL9A3) that contributed most to the classifications and were selected by the algorithms (random forest without RFE and with CV, CatBoost with and without RFE, XGBoost with and without RFE, and random forest with RFE and CV) (Fig. 6).

**Figure 6.**
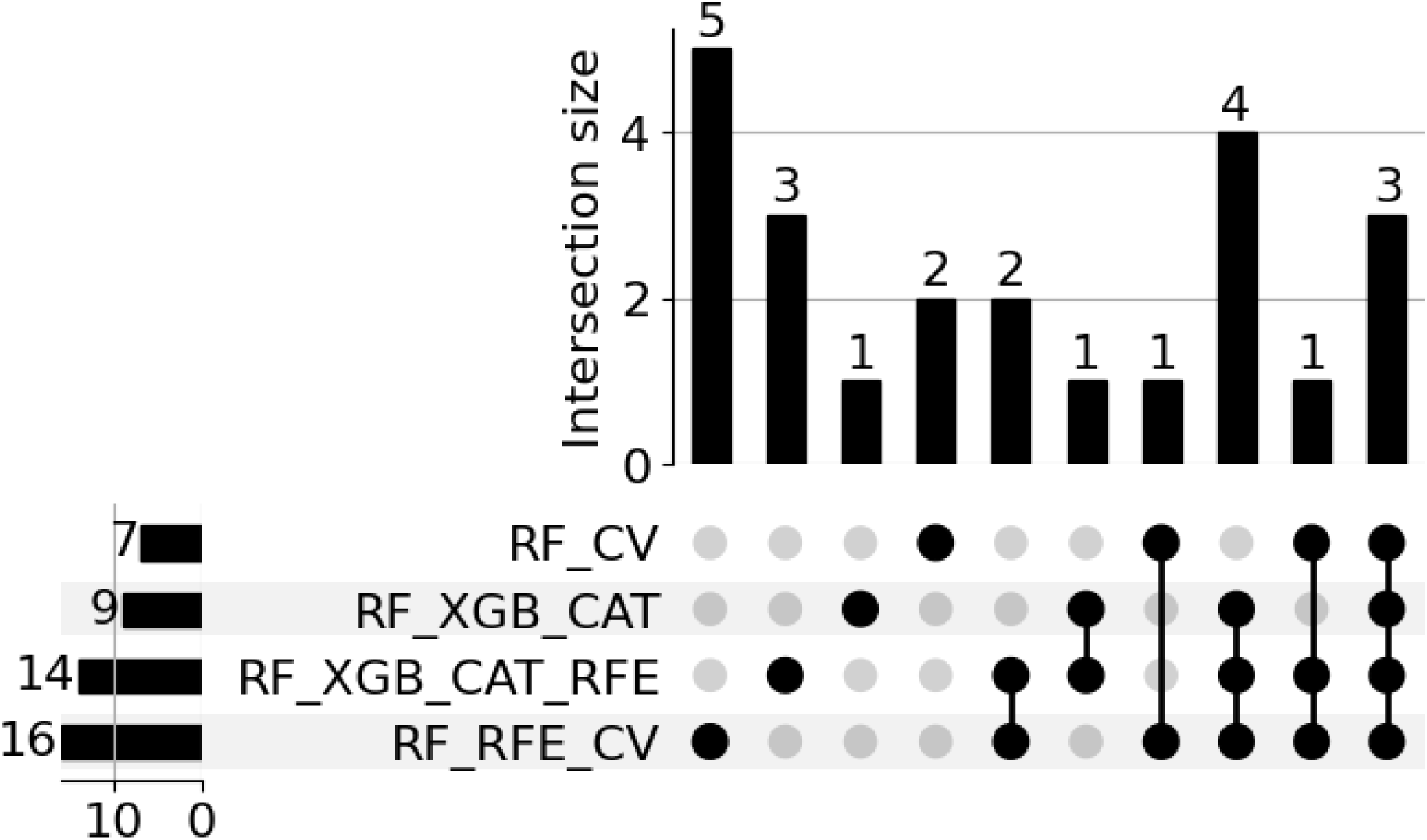
Upset plot with intersections of the best-tested models.

Based on these results, the 16 genes selected by all random forest folds may be good predictors of LUSC staging, especially when highlighting the 3 genes selected by all algorithms that performed well.

Next, we found some of the genes used in the post-RFE classification by observing genes present in pathways associated with lung cancer. The ESCO2 gene participates in the cell cycle regulation and is associated with cellular cohesion (Fig. 7). The COL9A3, MMP20, and GINS2 genes are associated with the PI3K ATK signaling pathway; COL9A3 acts in activating the PIK3CA gene, an oncogene which regulates important processes for carcinogenesis, such as cell proliferation, angiogenesis, DNA repair, cell survival, protein synthesis, and metabolism (Fig. 7). The CACNA1I gene is one of those responsible for the activation of the RAS family genes, inducing proliferation and differentiation in several types of cancer. Some genes are involved in associated processes, such as PLXNB3 via the axon guidance pathway, COL9A3 via the focal adhesion pathway, and the C7 gene, which induces regulation of the actin cytoskeleton. The GCGR and ALDH3B1 genes are associated with carbohydrate metabolism pathways, while ALDH3B1 and FTCD act in the histidine metabolism pathway. TMPRSS3 is a gene associated with Influenza A. The ESCO2 and PCAT7 genes act in the cell cycle pathway, and the H2AC14 gene acts in necroptosis.

**Figure 7.**
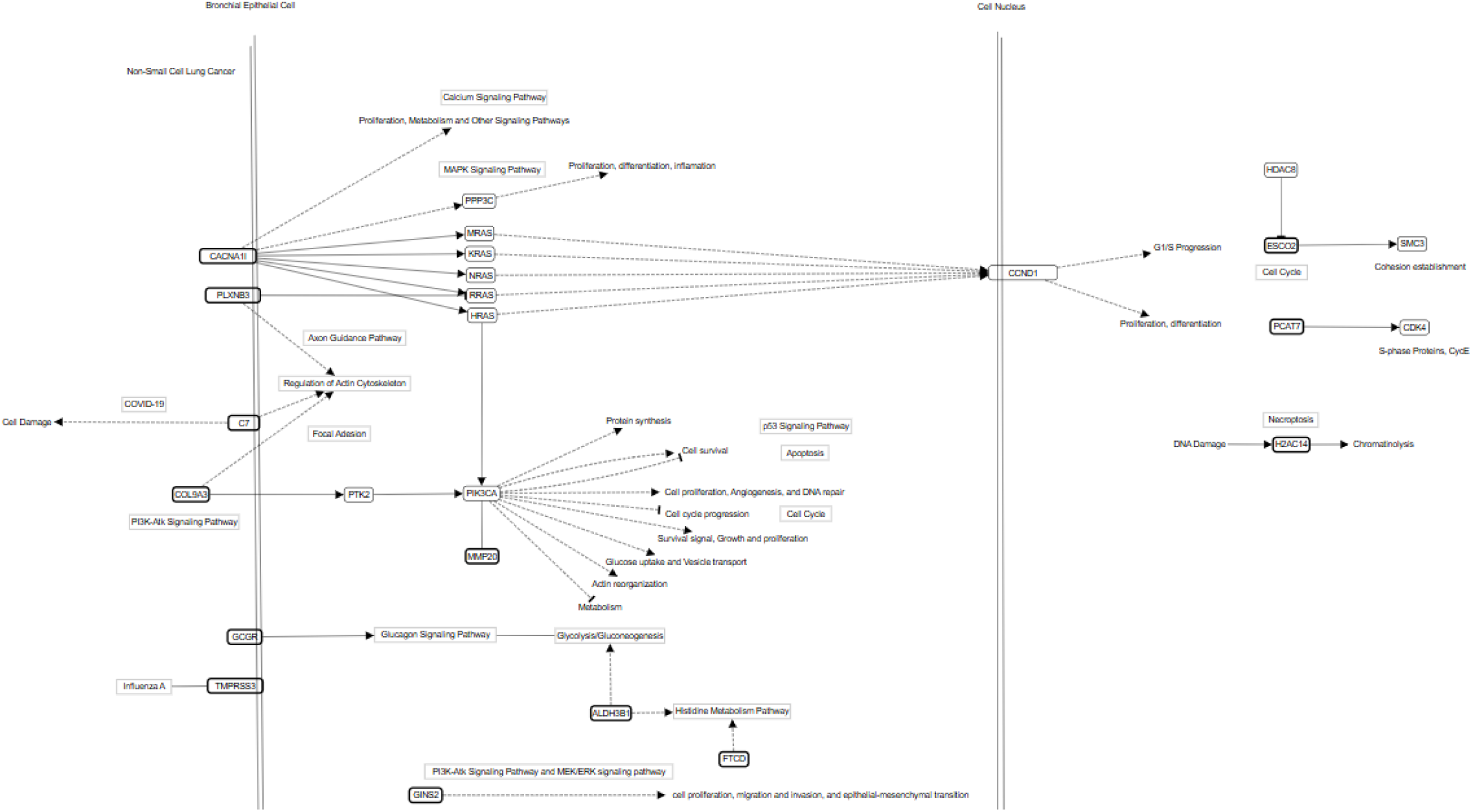
Pathway members and interactions are summarized in pathways. Rectangles with rounded edges indicate genes, and items with bold edges indicate genes present in the signature. Rectangles with gray borders represent pathways, and items outside rectangles indicate processes involving genes.

## Discussion

### Preprocessing, Machine Learning, and Classification

The 87 genes initially used for classifications came from a previous study [29]. The genes in this study were selected based on survival classifiers for different TCGA-LUSC subtypes. We observed the survival curve of the early and advanced subgroups to verify whether these 87 genes could be correlated with staging (Supplementary Fig. 1). Since survival and staging are correlated, we continued with the 87 genes. Then, we used RFE to obtain a smaller and more accurate signature, and it was possible to obtain classifications with more than 0.9 accuracy for the 38 selected genes (Table 3).

In addition to RFE and cross-validation, we also balanced the dataset, as very poor metrics were initially obtained when testing the classifiers with unbalanced data; all algorithms were classifying all patients as early stage (majority class). According to the average F1 score, the best balancing method was oversampling (Table 3), especially SMOTE. This technique was essential for obtaining the gene signature. It is widely used for many health problems and was associated with XAI to study cardiovascular risk assessment [44]. Good results were also found in studies on stroke prediction with different machine learning methods [45], combined with random forest in studies of metabolic syndrome [46], and cervical cancer [47].

After implementing RFE, CV, and data balancing, we observed that the best methods were three ensemble methods: random forest, CatBoost, and XGBoost. Random forest was the best model considering the area under the curve (Fig. 2) and accuracy, F1-score, precision, sensitivity, and specificity (Table 4). Random forest has been commonly used for this type of problem [29], [48], [49], [50], [51] because it is a robust and interpretable model, being able to rank variables [51], [52].

### Explainable Artificial Intelligence and Gene Signature

Given the complexity of the molecular mechanisms underlying LUSC and the challenges associated with predictive modeling for biomarker discovery, explainable artificial intelligence (XAI) emerges as a valuable strategy. Recent research in the field of lung cancer increasingly integrates machine learning with XAI, leveraging clinical, imaging, and molecular data to enhance predictive accuracy and interpretability. However, much of the work developed is directed to non-small cell lung cancers in general [21], [22], [23], [53]. Molecular therapies targeting LUSC are challenging due to their heterogeneity and complex genomic profile. Current molecular treatments based on PD-L1 are used non-specifically for non-small cell lung cancers. Nevertheless, this biomarker is less effective in LUSC than in LUAD [5]. Therefore, there is a need to search for other biomarkers specific to LUSC.

Furthermore, in order to compare potential LUSC gene signatures and their performance for staging classification, we checked different literature gene signatures associated with necroptosis [77], inflammation-based risk stratification [78], immune-related signatures [80], [43], aging [81] and EMT-associated genes [82], as well as prognostic [75], [79], [83] and cell death-related markers [76], [84]. These signatures reflect diverse biological processes and support prognosis and treatment response prediction in lung squamous cell carcinoma. Only the gene MMP20 from our signature was present in a prior study [43].

From this analysis we observed that 38 and 20 gene signatures obtained by our research reached the best performances according to the F1-score in all tested algorithms; in addition, the algorithms with the best performances for all tested signatures were CatBoost, random forest and XGBoost, which is in agreement with the results obtained by our signature.

Some genes with potential were observed in this search for new biomarkers associated with staging. The genes selected by all algorithms (Fig. 5) were MYOSLID, IMPDH1P8, and COL9A3. MYOSLID is a LncRNA associated with regulation of smooth muscle differentiation, and with the progression of head and neck squamous cell carcinoma and gastric cancer. Upregulation of this gene in osteosarcoma is associated with a poor prognosis, while its suppression reduces cell proliferation, migration, and invasion [54]. High IMPDH1P8 expression has an impact on the classification for advanced staging (Fig. 3), and high IMPDH1 expression is associated with poor prognosis in several tumors. In addition, this gene is associated with immune cells and immune checkpoints and is a potential prognostic marker and therapeutic target [55]. Then, low COL9A3 expression is associated with advanced stages of LUSC (Fig. 3). COL9A3 acts as a tumor suppressor in rectal cancer, is associated with prognosis in triple-negative breast cancer, and is an immune-related gene and a discriminator of good prognosis in esophageal squamous cell carcinoma [56]. Regarding its action in signaling pathways, COL9A3 is a gene associated with focal adhesion; it induces activation of the PIK3CA gene present in the PI3K-AKT pathway which in turn induces several important processes such as cell proliferation, angiogenesis, DNA repair, cell survival, protein synthesis, and metabolism. The presence of these components in the signature is encouraging, and it may be advantageous for immunotherapies, especially when combined with targeted treatments [5].

In addition to MYOSLID, IMPDH1P8, and COL9A3, other genes were selected by all random forest folds (Fig. 5). High MMP20, PCAT7, ALG1L, HMGB3P6, PLAAT1, GINS2, C7, ONECUT3, and ESCO2 gene expressions impact the classification for advanced staging. The GKN2 gene is associated with immune modulation and tumorigenesis and is considerably down-regulated in gastric and non-small cell lung cancers, having potential as a clinical biomarker [57], [58].

MMPs are matrix metalloproteinases which play a significant role in tissue organization, cellular differentiation, immune response, and carcinogenesis [59]. MMP20 plays a role in lung development and repair; its high expression is associated with tumorigenesis, and its high secretion is associated with metastatic activity. In addition, MMP20 regulates PI3-K and possibly c-myc activities, impacting signaling pathways that are vital for developing and maintaining lung tissue [60].

PCAT7 is a lncRNA whose expression in non-small cell lung cancers is high compared to normal cells, and its silencing considerably inhibits the progression of tumor cells. This gene is associated with the cell cycle pathway in lung cancer via miR-486-5p/CDK4 [61]. Moreover, high expression and hypomethylation levels of the ALG1L gene are associated with poor prognosis in lung cancer patients and have potential as a druggable target [62].

PLAAT1 has a negative correlation with the survival of LUSC patients. According to this study, GCGR present in 3 of the random forest folds has this same correlation [63]. In addition, PLAAT1 and ONECUT3 are immune-related genes associated with the prognosis in patients with LUSC [64]. ONECUT3 plays a role in cell differentiation and metabolism [65], and patients with high expression levels have poor survival [66].

In turn, high GINS2 expression is characteristic of several types of aggressive tumors [67] and promotes cell proliferation, migration, invasion, and epithelial-mesenchymal transition through the PI3K/Akt and MEK/ERK signaling pathways [68]. Then, C7 is a possible prognostic marker and target for immunotherapy in prostate cancer [69]. High ESCO2 expression is common in lung cancer, promotes cell proliferation and migration, and may promote malignancy of lung cancer cells through interference with the p53/p21 pathway [70]. Furthermore, ESCO2 also regulates the cell cycle pathway, inducing cellular cohesion (Fig. 7).

The CSTL1, H2AC14, and LINC02830 genes have low expression, which impacts the classification for advanced staging. The CSTL1 gene has already been cited in a signature associated with survival in lung cancer [71] and prognosis in colorectal cancer [72]. The H2AC14 gene correlates with poor prognosis of esophageal adenocarcinoma; however, its role in tumor development is not well elucidated [73]. According to KEGG [36], [37], [38], the H2AC14 gene is in the necroptosis pathway and induces chromatinolysis through DNA damage. In addition, the relationship between the HMGB3P6 and LINC02830 genes and the development of LUSC also needs to be better studied.

Some genes were not selected in all folds but were used for classification with RF, and some of these genes may have important biological roles. The ALDH3B1 and FTCD genes participate in the histidine metabolism pathway, and ALDH3B1 is present in the glycolysis/gluconeogenesis pathway along with GCGR.

The TMPRSS3 gene is in the influenza A pathway and is associated with alveolar and bronchial epithelial cells, alveolar macrophages, and type 2 pneumocytes [36], [37], [38]. The PLXNB3 gene is in the axon guidance pathway, which is associated with the actin cytoskeleton regulation pathway; the C7 gene induces focal adhesion processes in this pathway [36], [37], [38]. Regulation of the actin cytoskeleton may play a role in the epithelial-to-mesenchymal transition process in NSCLC.

The C7 gene induces activation of the Rho gene in regulating the actin cytoskeleton and consequently the formation of stress fibers. Rho activation may have a significant influence on cell morphology, as well as polarity and regulation of the metastatic spread of cancer cells [74].

The CACNA1I gene is in the calcium signaling pathway and induces proliferation, fertilization, learning and memory, contraction, metabolism, apoptosis, and exocytosis secretion. It is also in the MAPK signaling pathway, inducing proliferation and differentiation (Fig. 7).

The biological processes involving the signature genes are highly complex, and their specific roles in the progression of LUSC require further elucidation. However, as shown in Fig. 7, key pathways associated with LUSC are linked to genes within the signature, some of which are involved in driving various carcinogenic processes.

Many of the genes included in this signature are linked to important biological processes, including cell cycle regulation, tumorigenesis, cell differentiation, metastasis, cell migration, and immune response. Additionally, some of these genes (IMPDH1, GKN2, C7, and ALG1L) have already been identified as potential druggable targets [55], [58], [62], [69].

In future work, we aim to incorporate additional molecular data, such as copy number alterations and methylation profiles. These data may complement our knowledge of the biological processes associated with LUSC and help to integrate insights using external datasets.

## Conclusion

Given the complex relationship between gene expression and cancer progression, we evaluated multiple algorithms and data-balancing techniques. Notably, random forest and CatBoost achieved outstanding performance, reaching 90% accuracy.

Our findings indicate that the 38 selected genes play a crucial role in cancer staging. Furthermore, integrating machine learning with explainable AI (XAI) demonstrates significant potential in cancer research, providing a valuable tool for identifying staging marker genes in lung squamous cell carcinoma. XAI techniques enabled identifying the most influential variables in classification models, highlighting a potential gene signature consisting of the 20 genes selected by random forest. Among these, 16 genes were consistently selected across all random forest folds, while 3 genes were identified by SHAP in every test conducted with random forest, CatBoost, and XGBoost.

Many of the selected genes are associated with immune cells and immune checkpoints (IMPDH1, COL9A3, GKN2, PLAAT1, ONECUT3, and C7), suggesting their potential as immunotherapy targets. Although the collective role of these genes in developing lung squamous cell carcinoma (LUSC) remains unclear, several have been linked to other cancer types and key biological processes such as cell cycle regulation, proliferation, migration, and differentiation. This suggests that their expression may be relevant to LUSC carcinogenesis and that they could serve as potential targets for molecular therapy.

## Fundings

This work was supported in part by a grant number 88887.600025/2021-00 of Brazilian Funding agency CAPES-National Coordination of High Education Personnel Formation Programs. This work was supported in part by a grant number 88887.712333/2022-00 of Brazilian Funding agency CAPES-National Coordination of High Education Personnel Formation Programs. The article processing charge was funded by the Federal University of Rio Grande do Norte.

## Acknowledgment

The authors thank the Center for High-Performance Computing (Núcleo de Processamento de Alto Desempenho - NPAD/UFRN) available at https://npad.ufrn.br, the Multidisciplinary Bioinformatics Environment (BioME) at UFRN for providing computing resources for data processing, and the Brazilian Funding agencies CAPES and CNPq.

## Supplemental Figures and Tables

**Figure S1.**
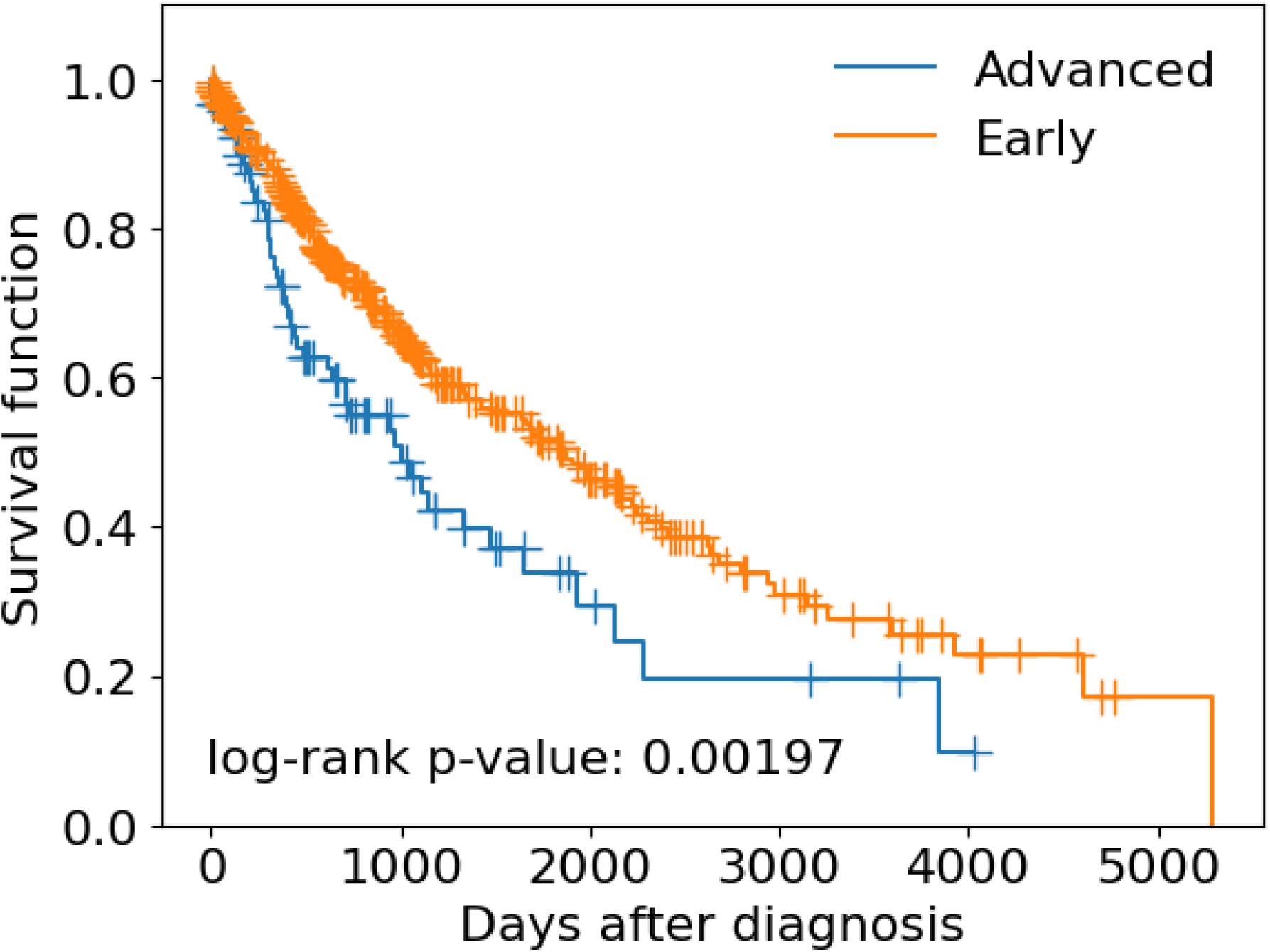
Survival Curves for Advanced and Early stages.

**Supplementary Table 1.**
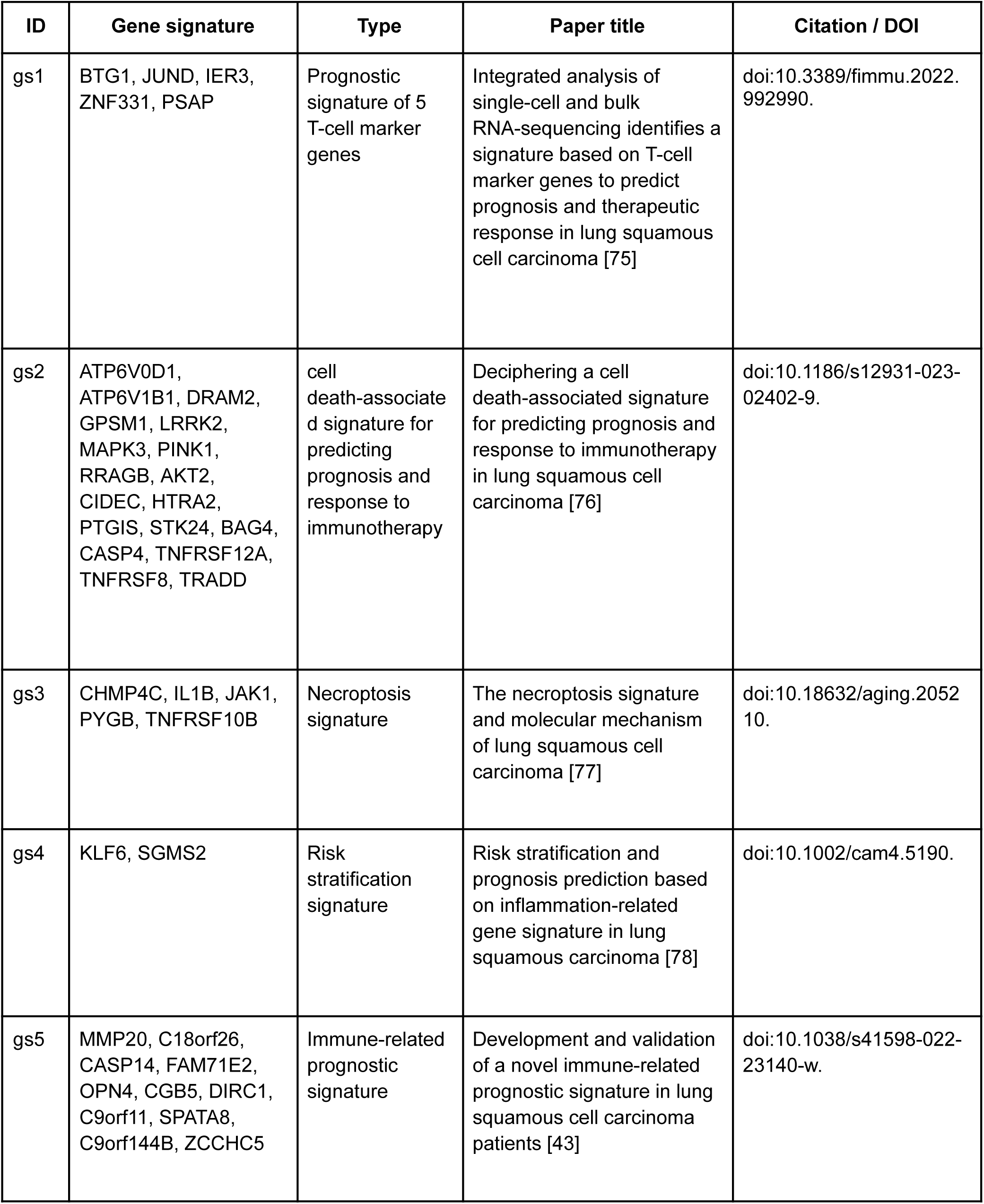
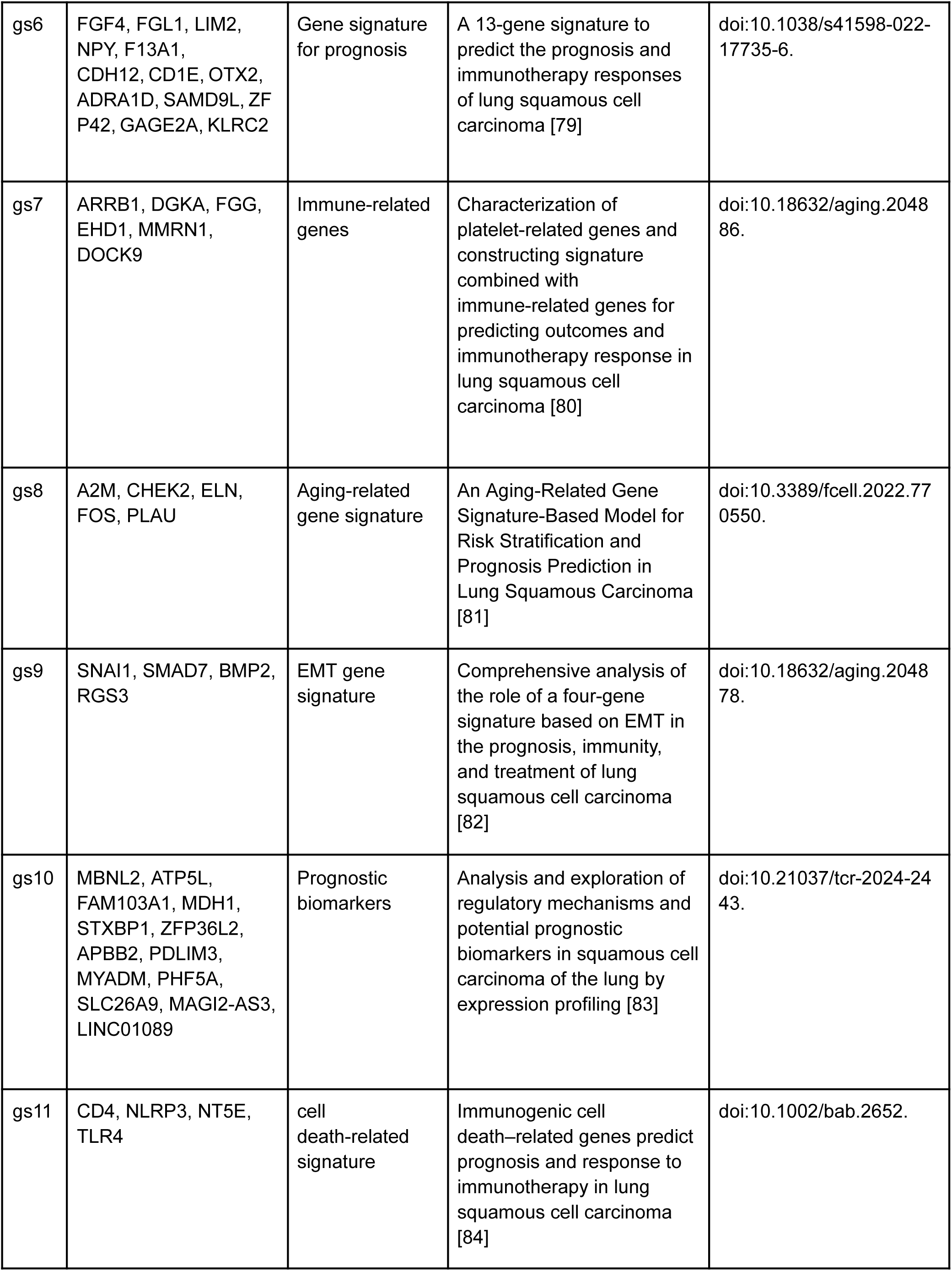
Gene signatures previously reported in eleven gene signatures on the literature reported in recent LUSC studies.

## Notes

### Competing Interest Statement

The authors have declared no competing interest.

## References

[1] M. B. Schabath and M. L. Cote, “Cancer Progress and Priorities: Lung Cancer,” Cancer Epidemiol. Biomarkers Prev., vol. 28, no. 10, pp. 1563–1579, Oct. 2019, doi: 10.1158/1055-9965.EPI-19-0221.

[2] R. L. Siegel, K. D. Miller, N. S. Wagle, and A. Jemal, “Cancer statistics, 2023,” CA. Cancer J. Clin., vol. 73, no. 1, pp. 17–48, Jan. 2023, doi: 10.3322/caac.21763.

[3] P. Perez-Moreno, E. Brambilla, R. Thomas, and J.-C. Soria, “Squamous Cell Carcinoma of the Lung: Molecular Subtypes and Therapeutic Opportunities,” Clin. Cancer Res., vol. 18, no. 9, pp. 2443–2451, May 2012, doi: 10.1158/1078-0432.CCR-11-2370.

[4] D. R. Gandara, P. S. Hammerman, M. L. Sos, P. N. Lara, and F. R. Hirsch, “Squamous Cell Lung Cancer: From Tumor Genomics to Cancer Therapeutics,” Clin. Cancer Res., vol. 21, no. 10, pp. 2236–2243, May 2015, doi: 10.1158/1078-0432.CCR-14-3039.

[5] S. C. M. Lau, P. Yuanwang, V. Velcheti, and K. K. Wong, “Squamous cell lung cancer: Current landscape and future therapeutic options,” Cancer Cell, vol. 40, pp. 1279–1293, 2022, doi: 10.1016/j.ccell.2022.09.018.

[6] The Cancer Genome Atlas Research Network, “Comprehensive genomic characterization of squamous cell lung cancers,” Nature, vol. 489, no. 7417, pp. 519–525, Sep. 2012, doi: 10.1038/nature11404.

[7] J. C. Smith and J. M. Sheltzer, “Genome-wide identification and analysis of prognostic features in human cancers,” Cell Rep., vol. 38, no. 13, p. 110569, Mar. 2022, doi: 10.1016/j.celrep.2022.110569.

[8] S. Dash, S. K. Shakyawar, M. Sharma, and S. Kaushik, “Big data in healthcare: management, analysis and future prospects,” J. Big Data, vol. 6, no. 1, p. 54, Dec. 2019, doi: 10.1186/s40537-019-0217-0.

[9] A. Dhillon and A. Singh, “Machine Learning in Healthcare Data Analysis: A Survey,” J. Biol. Todays World, vol. 8, p. 10, 2019.

[10] W. Zhu, L. Xie, J. Han, and X. Guo, “The Application of Deep Learning in Cancer Prognosis Prediction,” Cancers, vol. 12, no. 3, p. 603, Mar. 2020, doi: 10.3390/cancers12030603.

[11] T. Ching, X. Zhu, and L. X. Garmire, “Cox-nnet: An artificial neural network method for prognosis prediction of high-throughput omics data,” PLOS Comput. Biol., vol. 14, no. 4, p. e1006076, Apr. 2018, doi: 10.1371/journal.pcbi.1006076.

[12] M. Bakator and D. Radosav, “Deep Learning and Medical Diagnosis: A Review of Literature,” Multimodal Technol. Interact., vol. 2, no. 3, p. 47, Aug. 2018, doi: 10.3390/mti2030047.

[13] M. Gao, W. Kong, Z. Huang, and Z. Xie, “Identification of Key Genes Related to Lung Squamous Cell Carcinoma Using Bioinformatics Analysis,” Int. J. Mol. Sci., vol. 21, no. 8, p. 2994, Apr. 2020, doi: 10.3390/ijms21082994.

[14] K. Swanson, E. Wu, A. Zhang, A. A. Alizadeh, and J. Zou, “From patterns to patients: Advances in clinical machine learning for cancer diagnosis, prognosis, and treatment,” Cell, vol. 186, no. 8, pp. 1772–1791, Apr. 2023, doi: 10.1016/j.cell.2023.01.035.

[15] P. Terrematte, D. Andrade, J. Justino, B. Stransky, D. de Araújo, and A. Dória Neto, “A Novel Machine Learning 13-Gene Signature: Improving Risk Analysis and Survival Prediction for Clear Cell Renal Cell Carcinoma Patients,” Cancers, vol. 14, no. 9, p. 2111, Apr. 2022, doi: 10.3390/cancers14092111.

[16] M. R. Karim et al., “Explainable AI for Bioinformatics: Methods, Tools and Applications,” Brief Bioinform, vol. 24, no. 5, 2023, doi: 10.1093/bib/bbad236.

[17] C. Ladbury et al., “Utilization of model-agnostic explainable artificial intelligence frameworks in oncology: a narrative review,” Transl. Cancer Res., vol. 11, no. 10, pp. 3853–3868, Oct. 2022, doi: 10.21037/tcr-22-1626.

[18] R. Yang, X. Xiong, H. Wang, and W. Li, “Explainable Machine Learning Model to Prediction EGFR Mutation in Lung Cancer,” Front. Oncol., vol. 12, p. 924144, 2022, doi: 10.3389/fonc.2022.924144.

[19] B. Alsinglawi et al., “An explainable machine learning framework for lung cancer hospital length of stay prediction,” Sci. Rep., vol. 12, no. 1, p. 607, Jan. 2022, doi: 10.1038/s41598-021-04608-7.

[20] S. T. Rikta, K. M. M. Uddin, N. Biswas, R. Mostafiz, F. Sharmin, and S. K. Dey, “XML-GBM lung: An explainable machine learning-based application for the diagnosis of lung cancer,” J. Pathol. Inform., vol. 14, p. 100307, 2023, doi: 10.1016/j.jpi.2023.100307.

[21] K. Dwivedi, A. Rajpal, S. Rajpal, M. Agarwal, V. Kumar, and N. Kumar, “An explainable AI-driven biomarker discovery framework for Non-Small Cell Lung Cancer classification,” Comput. Biol. Med., vol. 153, p. 106544, Feb. 2023, doi: 10.1016/j.compbiomed.2023.106544.

[22] C. Ladbury et al., “Explainable Artificial Intelligence to Identify Dosimetric Predictors of Toxicity in Patients with Locally Advanced Non-Small Cell Lung Cancer: A Secondary Analysis of RTOG 0617,” Int. J. Radiat. Oncol. Biol. Phys., vol. 117, no. 5, pp. 1287–1296, Dec. 2023, doi: 10.1016/j.ijrobp.2023.06.019.

[23] J. R. Astley, J. M. Reilly, S. Robinson, J. M. Wild, M. Q. Hatton, and B. A. Tahir, “Explainable deep learning-based survival prediction for non-small cell lung cancer patients undergoing radical radiotherapy,” Radiother. Oncol., vol. 193, p. 110084, Apr. 2024, doi: 10.1016/j.radonc.2024.110084.

[24] “GDC,” GDC Data Portal. Accessed: Mar. 21, 2024. [Online]. Available: https://portal.gdc.cancer.gov/

[25] A. Colaprico et al., “TCGAbiolinks: an R/Bioconductor package for integrative analysis of TCGA data,” Nucleic Acids Res., vol. 44, no. 8, pp. e71–e71, May 2016, doi: 10.1093/nar/gkv1507.

[26] “Xenabrowser.” Accessed: Mar. 21, 2024. [Online]. Available: https://xenabrowser.net/datapages/?cohort=TCGA%20Lung%20Squamous%20Cell%20Carcinoma%20(LUSC)&removeHub=https%3A%2F%2Fxena.treehouse.gi.ucsc.edu%3A443

[27] The pandas development team, pandas-dev/pandas: Pandas. (Sep. 20, 2024). Zenodo. doi: 10.5281/ZENODO.3509134.

[28] C. Davidson-Pilon, lifelines, survival analysis in Python. (Oct. 29, 2024). Zenodo. doi: 10.5281/ZENODO.805993.

[29] D. V. C. Lima, P. Terrematte, B. Stransky, and A. D. D. Neto, “An Integrated Data Analysis Using Bioinformatics and Random Forest to Predict Prognosis of Patients With Squamous Cell Lung Cancer,” IEEE Access, vol. 12, pp. 59335–59345, 2024, doi: 10.1109/ACCESS.2024.3392277.

[30] F. Pedregosa et al., “Scikit-learn: Machine Learning in Python,” J. Mach. Learn. Res., vol. 12, no. 85, pp. 2825–2830, 2011.

[31] G. Lema, F. Nogueira, and C. K. Aridas, “Imbalanced-learn: A Python Toolbox to Tackle the Curse of Imbalanced Datasets in Machine Learning,” JMLR, vol. 18, no. 17, pp. 1–5, 2017.

[32] A. V. Dorogush, V. Ershov, and A. Gulin, “CatBoost: gradient boosting with categorical features support,” 2018, doi: 10.48550/ARXIV.1810.11363.

[33] J. D. Hunter, “Matplotlib: A 2D graphics environment,” Computing in Science \& Engineering, vol. 9, no. 3, pp. 90–95, 2007, doi: 10.1109/MCSE.2007.55.

[34] S. M. Lundberg and S.-I. Lee, “A Unified Approach to Interpreting Model Predictions,” in Advances in Neural Information Processing Systems, I. Guyon, U. V. Luxburg, S. Bengio, H. Wallach, R. Fergus, S. Vishwanathan, and R. Garnett, Eds., Curran Associates, Inc., 2017. [Online]. Available: https://proceedings.neurips.cc/paper_files/paper/2017/file/8a20a8621978632d76c43dfd28b67767-Paper.pdf

[35] A. Lex, N. Gehlenborg, H. Strobelt, R. Vuillemot, and H. Pfister, “UpSet: Visualization of Intersecting Sets,” IEEE Trans. Vis. Comput. Graph., vol. 20, no. 12, pp. 1983–1992, Dec. 2014, doi: 10.1109/TVCG.2014.2346248.

[36] M. Kanehisa, “KEGG: Kyoto Encyclopedia of Genes and Genomes,” Nucleic Acids Res., vol. 28, no. 1, pp. 27–30, Jan. 2000, doi: 10.1093/nar/28.1.27.

[37] M. Kanehisa, “Toward understanding the origin and evolution of cellular organisms,” Protein Sci., vol. 28, no. 11, pp. 1947–1951, Nov. 2019, doi: 10.1002/pro.3715.

[38] M. Kanehisa, M. Furumichi, Y. Sato, Y. Matsuura, and M. Ishiguro-Watanabe, “KEGG: biological systems database as a model of the real world,” Nucleic Acids Res., vol. 53, no. D1, pp. D672–D677, Jan. 2025, doi: 10.1093/nar/gkae909.

[39] I. Bahceci, U. Dogrusoz, K. C. La, Ö. Babur, J. Gao, and N. Schultz, “PathwayMapper: a collaborative visual web editor for cancer pathways and genomic data,” Bioinformatics, vol. 33, no. 14, pp. 2238–2240, Jul. 2017, doi: 10.1093/bioinformatics/btx149.

[40] G. van Rossum and F. L. Drake, The Python language reference, Release 3.0.1 [Repr.]. in Python documentation manual / Guido van Rossum; Fred L. Drake [ed.], no. Pt. 2. Hampton, NH: Python Software Foundation, 2010.

[41] R. Ihaka and R. Gentleman, “R: A Language for Data Analysis and Graphics,” J. Comput. Graph. Stat., vol. 5, no. 3, pp. 299–314, Sep. 1996, doi: 10.1080/10618600.1996.10474713.

[42] RStudio Team, “RStudio | Open source & professional software for data science teams.” Accessed: Apr. 06, 2022. [Online]. Available: https://www.rstudio.com/

[43] X. Liu et al., “Development and validation of a novel immune-related prognostic signature in lung squamous cell carcinoma patients,” Sci. Rep., vol. 12, no. 1, p. 20737, Dec. 2022, doi: 10.1038/s41598-022-23140-w.

[44] F. Yang, Y. Qiao, P. Hajek, and M. Z. Abedin, “Enhancing cardiovascular risk assessment with advanced data balancing and domain knowledge-driven explainability,” Expert Syst. Appl., vol. 255, p. 124886, Dec. 2024, doi: 10.1016/j.eswa.2024.124886.

[45] Y. Wu and Y. Fang, “Stroke Prediction with Machine Learning Methods among Older Chinese,” Int. J. Environ. Res. Public. Health, vol. 17, no. 6, p. 1828, Mar. 2020, doi: 10.3390/ijerph17061828.

[46] S. Mohseni-Takalloo, H. Mohseni, H. Mozaffari-Khosravi, M. Mirzaei, and M. Hosseinzadeh, “The effect of data balancing approaches on the prediction of metabolic syndrome using non-invasive parameters based on random forest,” BMC Bioinformatics, vol. 25, no. 1, p. 18, Jan. 2024, doi: 10.1186/s12859-024-05633-9.

[47] M. Arora, S. Dhawan, and K. Singh, “Data Driven Prognosis of Cervical Cancer Using Class Balancing and Machine Learning Techniques,” EAI Endorsed Trans. Energy Web, p. 164264, Jul. 2018, doi: 10.4108/eai.13-7-2018.164264.

[48] K. Lee, H. Jeong, S. Lee, and W.-K. Jeong, “CPEM: Accurate cancer type classification based on somatic alterations using an ensemble of a random forest and a deep neural network,” Sci. Rep., vol. 9, no. 1, p. 16927, Nov. 2019, doi: 10.1038/s41598-019-53034-3.

[49] O. Abdelwahab, N. Awad, M. Elserafy, and E. Badr, “A feature selection-based framework to identify biomarkers for cancer diagnosis: A focus on lung adenocarcinoma,” PLOS ONE, vol. 17, no. 9, p. e0269126, Sep. 2022, doi: 10.1371/journal.pone.0269126.

[50] X. Wu, B.-B. Denise, F. Zhan, and J. Zhang, “Determining Association between Lung Cancer Mortality Worldwide and Risk Factors Using Fuzzy Inference Modeling and Random Forest Modeling,” Int. J. Environ. Res. Public. Health, vol. 19, no. 21, p. 14161, Oct. 2022, doi: 10.3390/ijerph192114161.

[51] X. Chen and H. Ishwaran, “Random forests for genomic data analysis,” Genomics, vol. 99, no. 6, pp. 323–329, Jun. 2012, doi: 10.1016/j.ygeno.2012.04.003.

[52] L. Breiman, “Random Forests,” Mach. Learn., vol. 45, no. 1, pp. 5–32, 2001, doi: 10.1023/A:1010933404324.

[53] C. Duan et al., “GWO+RuleFit: rule-based explainable machine-learning combined with heuristics to predict mid-treatment FDG PET response to chemoradiation for locally advanced non-small cell lung cancer,” Phys. Med. Biol., vol. 69, no. 15, p. 155018, Aug. 2024, doi: 10.1088/1361-6560/ad6118.

[54] S. Yang, M. Chen, and C. Lin, “A Novel lncRNA MYOSLID/miR-1286/RAB13 Axis Plays a Critical Role in Osteosarcoma Progression,” Cancer Manag. Res., vol. Volume 11, pp. 10345–10351, Dec. 2019, doi: 10.2147/CMAR.S231376.

[55] C. Liu, W. Zhang, X. Zhou, and L. Liu, “IMPDH1, a prognostic biomarker and immunotherapy target that correlates with tumor immune microenvironment in pan-cancer and hepatocellular carcinoma,” Front. Immunol., vol. 13, p. 983490, Dec. 2022, doi: 10.3389/fimmu.2022.983490.

[56] M. Liu et al., “Profiles of immune cell infiltration and immune-related genes in the tumor microenvironment of esophageal squamous cell carcinoma,” BMC Med. Genomics, vol. 14, no. 1, p. 75, Dec. 2021, doi: 10.1186/s12920-021-00928-9.

[57] F. Liu and H. Wu, “Prognostic Value of Gastrokine-2 (GKN2) and Its Correlation with Tumor-Infiltrating Immune Cells in Lung Cancer and Gastric Cancers,” J. Inflamm. Res., vol. Volume 13, pp. 933–944, Nov. 2020, doi: 10.2147/JIR.S277353.

[58] X. Yang et al., “Low-level gastrokine 2 promoted progress of NSCLC and as a potential biomarker,” J.Clin. Lab. Anal., vol. 36, no. 2, p. e24213, Feb. 2022, doi: 10.1002/jcla.24213.

[59] C. Wei, “The multifaceted roles of matrix metalloproteinases in lung cancer,” Front. Oncol., vol. 13, p. 1195426, Sep. 2023, doi: 10.3389/fonc.2023.1195426.

[60] P. L. Mok et al., “Lung development, repair and cancer: A study on the role of MMP20 gene in adenocarcinoma,” PLOS ONE, vol. 16, no. 4, p. e0250552, Apr. 2021, doi: 10.1371/journal.pone.0250552.

[61] W. Geng, M. Qiu, D. Zhang, P. Li, G. Sun, and X. Zhou, “LncRNA PCAT7 promotes non-small cell lung cancer progression by activating miR-486-5p/CDK4 axis-mediated cell cycle,” Am. J. Transl. Res., vol. 14, no. 5, pp. 3003–3016, 2022.

[62] P. Han, Q. Liu, and J. Xiang, “Monitoring methylation-driven genes as prognostic biomarkers in patients with lung squamous cell cancer,” Oncol. Lett., Nov. 2019, doi: 10.3892/ol.2019.11163.

[63] H. Zhang, “Lung Squamous Cell Carcinoma (LUSC) Survival Analysis using TCGA Database,” in Proceedings of the 2023 4th International Symposium on Artificial Intelligence for Medicine Science, Chengdu China: ACM, Oct. 2023, pp. 1119–1129. doi: 10.1145/3644116.3644307.

[64] J. Pu et al., “Construction of a prognostic model for lung squamous cell carcinoma based on immune-related genes,” Carcinogenesis, vol. 44, no. 2, pp. 143–152, May 2023, doi: 10.1093/carcin/bgac098.

[65] K. Sunita Prajapati, S. Gupta, S. Chaudhri, and S. Kumar, “Role of ONECUT family transcription factors in cancer and other diseases,” Exp. Cell Res., vol. 438, no. 1, p. 114035, May 2024, doi: 10.1016/j.yexcr.2024.114035.

[66] H. Shi, Y. Tsang, Y. Yang, and H. L. Chin, “Identification of ONECUT3 as a stemness-related transcription factor regulating NK cell-mediated immune evasion in pancreatic cancer,” Sci. Rep., vol. 13, no. 1, p. 18133, Oct. 2023, doi: 10.1038/s41598-023-45560-y.

[67] D. Sun, Y. Zong, J. Cheng, Z. Li, L. Xing, and J. Yu, “GINS2 attenuates the development of lung cancer by inhibiting the STAT signaling pathway,” J. Cancer, vol. 12, no. 1, pp. 99–110, 2021, doi: 10.7150/jca.46744.

[68] X. Liu, L. Sun, S. Zhang, S. Zhang, and W. Li, “GINS2 facilitates epithelial-to-mesenchymal transition in non-small-cell lung cancer through modulating PI3K/Akt and MEK/ERK signaling,” J. Cell. Physiol., vol. 235, no. 11, pp. 7747–7756, Nov. 2020, doi: 10.1002/jcp.29381.

[69] Z. Chen et al., “Complement C7 (C7), a Potential Tumor Suppressor, Is an Immune-Related Prognostic Biomarker in Prostate Cancer (PC),” Front. Oncol., vol. 10, p. 1532, Aug. 2020, doi: 10.3389/fonc.2020.01532.

[70] M. Liu, Y. Ma, and E. Wang, “ESCO2 Inhibits p53 Transcription and Promotes Proliferation and Migration of Lung Cancer Cells,” May 11, 2021. doi: 10.21203/rs.3.rs-506320/v1.

[71] T. Miao, L. Du, W. Xiao, B. Mao, Y. Wang, and J. Fu, “Identification of Survival-Associated Gene Signature in Lung Cancer Coexisting With COPD,” Front. Oncol., vol. 11, p. 600243, Mar. 2021, doi: 10.3389/fonc.2021.600243.

[72] J. Zou et al., “Multi-Omics Analysis of the Tumor Microenvironment in Liver Metastasis of Colorectal Cancer Identified FJX1 as a Novel Biomarker,” Front. Genet., vol. 13, p. 960954, Jul. 2022, doi: 10.3389/fgene.2022.960954.

[73] S. Zhang et al., “Development and Validation of a Prognostic Model for Esophageal Adenocarcinoma Based on Necroptosis-Related Genes,” Genes, vol. 13, no. 12, p. 2243, Nov. 2022, doi: 10.3390/genes13122243.

[74] M. Cooke, M. J. Baker, M. G. Kazanietz, and V. Casado-Medrano, “PKCε regulates Rho GTPases and actin cytoskeleton reorganization in non-small cell lung cancer cells,” Small GTPases, vol. 12, no. 3, pp. 202–208, May 2021, doi: 10.1080/21541248.2019.1684785.

[75] X. Shi et al., “Integrated analysis of single-cell and bulk RNA-sequencing identifies a signature based on T-cell marker genes to predict prognosis and therapeutic response in lung squamous cell carcinoma,” Front. Immunol., vol. 13, p. 992990, Oct. 2022, doi: 10.3389/fimmu.2022.992990.

[76] G. Mao et al., “Deciphering a cell death-associated signature for predicting prognosis and response to immunotherapy in lung squamous cell carcinoma,” Respir. Res., vol. 24, no. 1, p. 176, Jul. 2023, doi: 10.1186/s12931-023-02402-9.

[77] G.-Q. Song et al., “The necroptosis signature and molecular mechanism of lung squamous cell carcinoma,” Aging, vol. 15, no. 22, pp. 12907–12926, Nov. 2023, doi: 10.18632/aging.205210.

[78] W. Zhai et al., “Risk stratification and prognosis prediction based on inflammation-related gene signature in lung squamous carcinoma,” Cancer Med., vol. 12, no. 4, pp. 4968–4980, Feb. 2023, doi: 10.1002/cam4.5190.

[79] Q. Yang, H. Gong, J. Liu, M. Ye, W. Zou, and H. Li, “A 13-gene signature to predict the prognosis and immunotherapy responses of lung squamous cell carcinoma,” Sci. Rep., vol. 12, no. 1, p. 13646, Aug. 2022, doi: 10.1038/s41598-022-17735-6.

[80] S. Zhao, H. Gong, and W. Liang, “Characterization of platelet-related genes and constructing signature combined with immune-related genes for predicting outcomes and immunotherapy response in lung squamous cell carcinoma,” Aging, vol. 15, no. 14, pp. 6969–6992, Jul. 2023, doi: 10.18632/aging.204886.

[81] W.-Y. Zhai et al., “An Aging-Related Gene Signature-Based Model for Risk Stratification and Prognosis Prediction in Lung Squamous Carcinoma,” Front. Cell Dev. Biol., vol. 10, p. 770550, Mar. 2022, doi: 10.3389/fcell.2022.770550.

[82] F. Li et al., “Comprehensive analysis of the role of a four-gene signature based on EMT in the prognosis, immunity, and treatment of lung squamous cell carcinoma,” Aging, vol. 15, no. 14, pp. 6865–6893, Jul. 2023, doi: 10.18632/aging.204878.

[83] X. Wan et al., “Analysis and exploration of regulatory mechanisms and potential prognostic biomarkers in squamous cell carcinoma of the lung by expression profiling,” Transl. Cancer Res., vol. 14, no. 1, pp. 569–583, Jan. 2025, doi: 10.21037/tcr-2024-2443.

[84] G. Li, K. Chen, S. Dong, X. Wei, L. Zhou, and B. Wang, “Immunogenic cell death–related genes predict prognosis and response to immunotherapy in lung squamous cell carcinoma,” Biotechnol. Appl. Biochem., vol. 72, no. 1, pp. 138–149, Feb. 2025, doi: 10.1002/bab.2652.

