## Supplemental Table 1 for "Explainable AI and Multiclassifiers for Staging Biomarker Discovery in Lung Squamous Cell Carcinoma"

Supplementary Table 1 - Gene signatures previously reported in eleven gene signatures on the literature reported in recent LUSC studies.

| ID | Gene signature | Type | Paper title | Citation / DOI |
|---|---|---|---|---|
| gs1 | BTG1, JUND, IER3, ZNF331, PSAP | Prognostic signature of 5 T-cell marker genes | Integrated analysis of single-cell and bulk RNA-sequencing identifies a signature based on T-cell marker genes to predict prognosis and therapeutic response in lung squamous cell carcinoma [75] | doi:10.3389/fimmu.2022.992990. |
| gs2 | ATP6V0D1, ATP6V1B1, DRAM2, GPSM1, LRRK2, MAPK3, PINK1, RRAGB, AKT2, CIDEC, HTRA2, PTGIS, STK24, BAG4, CASP4, TNFRSF12A, TNFRSF8, TRADD | cell death-associated signature for predicting prognosis and response to immunotherapy | Deciphering a cell death-associated signature for predicting prognosis and response to immunotherapy in lung squamous cell carcinoma [76] | doi:10.1186/s12931-023-02402-9. |
| gs3 | CHMP4C, IL1B, JAK1, PYGB, TNFRSF10B | Necroptosis signature | The necroptosis signature and molecular mechanism of lung squamous cell carcinoma [77] | doi:10.18632/aging.205210. |
| gs4 | KLF6, SGMS2 | Risk stratification signature | Risk stratification and prognosis prediction based on inflammation-related gene signature in lung squamous carcinoma [78] | doi:10.1002/cam4.5190. |
| gs5 | MMP20, C18orf26, CASP14, FAM71E2, OPN4, CGB5, DIRC1, C9orf11, SPATA8, C9orf144B, ZCCHC5 | Immune-related prognostic signature | Development and validation of a novel immune-related prognostic signature in lung squamous cell carcinoma patients [43] | doi:10.1038/s41598-022-23140-w. |

| | | | | |
|---|---|---|---|---|
| gs6 | FGF4, FGL1, LIM2, NPY, F13A1, CDH12, CD1E, OTX2, ADRA1D, SAMD9L, ZFP42, GAGE2A, KLRC2 | Gene signature for prognosis | A 13-gene signature to predict the prognosis and immunotherapy responses of lung squamous cell carcinoma [79] | doi:10.1038/s41598-022-17735-6. |
| gs7 | ARRB1, DGKA, FGG, EHD1, MMRN1, DOCK9 | Immune-related genes | Characterization of platelet-related genes and constructing signature combined with immune-related genes for predicting outcomes and immunotherapy response in lung squamous cell carcinoma [80] | doi:10.18632/aging.204886. |
| gs8 | A2M, CHEK2, ELN, FOS, PLAU | Aging-related gene signature | An Aging-Related Gene Signature-Based Model for Risk Stratification and Prognosis Prediction in Lung Squamous Carcinoma [81] | doi:10.3389/fcell.2022.770550. |
| gs9 | SNAI1, SMAD7, BMP2, RGS3 | EMT gene signature | Comprehensive analysis of the role of a four-gene signature based on EMT in the prognosis, immunity, and treatment of lung squamous cell carcinoma [82] | doi:10.18632/aging.204878. |
| gs10 | MBNL2, ATP5L, FAM103A1, MDH1, STXBP1, ZFP36L2, APBB2, PDLIM3, MYADM, PHF5A, SLC26A9, MAGI2-AS3, LINC01089 | Prognostic biomarkers | Analysis and exploration of regulatory mechanisms and potential prognostic biomarkers in squamous cell carcinoma of the lung by expression profiling [83] | doi:10.21037/tcr-2024-2443. |
| gs11 | CD4, NLRP3, NT5E, TLR4 | cell death-related signature | Immunogenic cell death–related genes predict prognosis and response to immunotherapy in lung squamous cell carcinoma [84] | doi:10.1002/bab.2652. |
