## Supplementary figures and images for "Explainable AI and Multiclassifiers for Staging Biomarker Discovery in Lung Squamous Cell Carcinoma"

### Supplemental Figure 1

**Supplemental Figures and Tables**

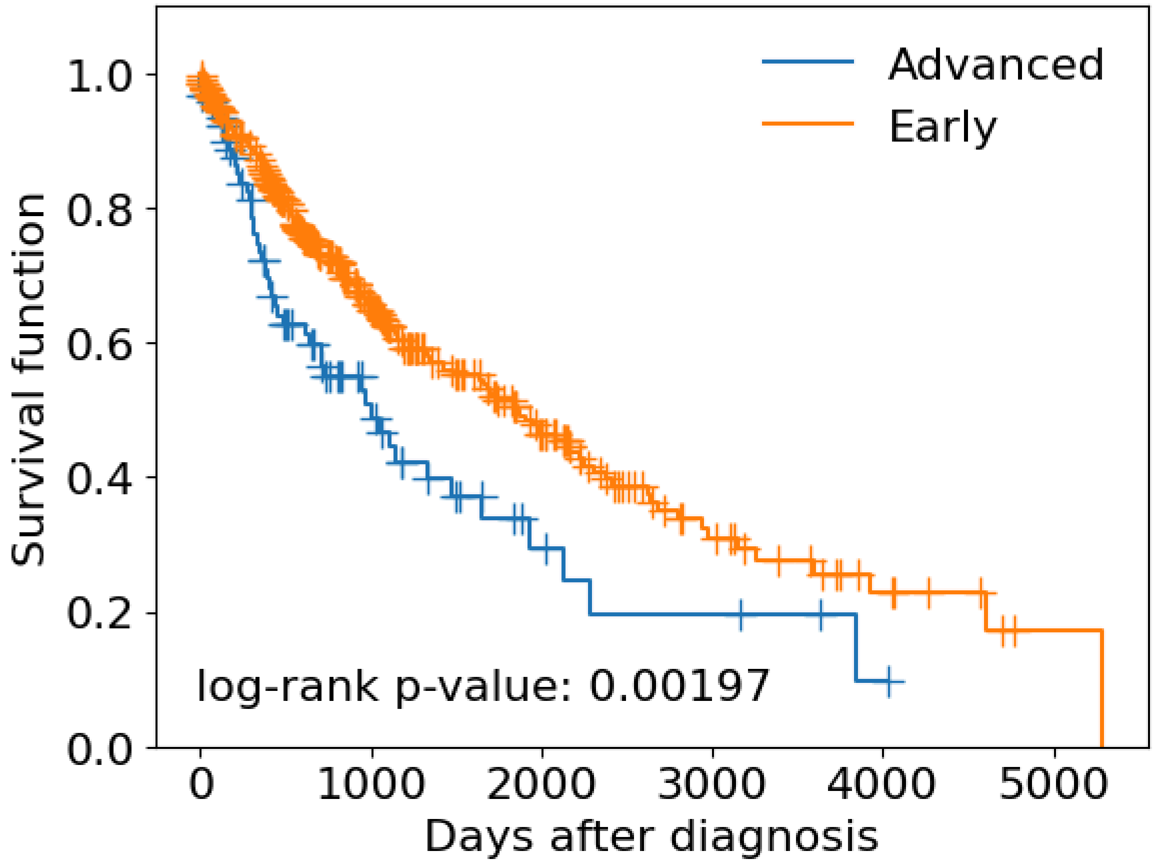


**Figure S1 - Survival Curves for Advanced and Early stages.**
